# Bacterial nanotubes extend from a tubeosome organelle to establish an intercellular bridge

**DOI:** 10.64898/2026.07.27.740770

**Authors:** Manas Kumar Guria, Tengfei Zhong, Amit K. Baidya, Małgorzata Zuzanna Zora, Ritesh Ranjan Pal, Venkadesaperumal Gopu, Saurabh Bhattacharya, Oren Yakovian, Miriam Ravins, Debnath Ghosal, Grant J. Jensen, Ilan Rosenshine, Mohammed Kaplan, Sigal Ben-Yehuda

**Author notes:** These authors contributed equally to this work (Manas Kumar Guria, Tengfei Zhong). Molecular Biophysics Unit, Division of Biological Sciences, Indian Institute of Science, Bangalore - 560012, India (Amit K. Baidya). School of Biological Sciences, Indian Association for the Cultivation of Science, Kolkata 700032, India (Ritesh Ranjan Pal). To whom correspondence should be addressed. (IR), (MK), (S. B-Y).

## Abstract

Bacterial nanotubes are widespread intercellular membranous bridges mediating molecular exchange among neighboring cells, yet their formation and dynamics remain largely unexplored. By combining live-cell imaging with *in situ* cryo-electron tomography, we describe nanotube biogenesis in the Gram-positive *Bacillus subtilis*. Nanotubes form elaborate, highly dynamic networks, with extending nanotubes exhibiting multidirectional movement toward nearby cells, repeatedly scanning their surface. Ultimately, nanotubes penetrate through the recipient cell wall to establish an intercellular bridge. Strikingly, nanotubes emerge within milliseconds from lipid-enriched sites representing a previously unrecognized membranous organelle, termed tubeosome, located in an unusual pseudo-periplasmic space beneath the cell wall. We present evidence that tubeosomes comprise proto-nanotube reservoirs that unfold to project nanotubes. Our study uncovers the tubeosome as a novel bacterial organelle and provides the first real-time visualization of intercellular bridge formation. As connecting nanotubes are found across bacteria, archaea, and eukaryotes, this mechanism may reflect a broadly conserved strategy for cellular communication.

## Introduction

Bacteria in natural habitats assemble into intricate multi-species communities typified by intimate contact-dependent interactions with neighbouring cells. These interactions often involve dedicated proteinaceous protrusions, such as type IV, VI, and VII secretion systems, exploited for intercellular molecular delivery [e.g., ^1-5^]. A less-explored mode of bacterial intercellular route is mediated by membranous conduits, termed nanotubes ^6,7^. Bacterial nanotubes were shown to be utilized for intercellular trafficking of cytoplasmic proteins and even small non-conjugative plasmids ^6-10^. Interestingly, we have recently discovered a family of nucleases that specifically restricts nanotube-mediated plasmid exchange, highlighting their potential role in horizontal gene transfer ^11^. Nanotube-like structures have additionally been linked to nutrient cross-feeding between bacteria of different species [e.g., ^10,12-14^], with recent studies demonstrating nanotube-mediated exchange among marine cyanobacteria, supporting their role both within and between genera ^15^. Networks of nanotubes have also been implicated in host-nutrient extraction by pathogenic and symbiotic bacteria ^16,17^.

In the Gram-positive bacterium *Bacillus subtilis* (*B. subtilis*), a calcineurin-like phosphodiesterase, YmdB, is localized to nanotubes and facilitates their formation ^8,18^, whereas the cell wall remodelling enzymes, LytC amidase and its enhancer LytB, are required for efficient nanotube extrusion and penetration into recipient bacteria ^19^. Furthermore, a membrane-associated complex of five conserved proteins, FliP, FliQ, FliR, FlhB, and FlhA, designated CORE, constituting the flagellar export apparatus, was found to be key for nanotube biogenesis. CORE-dependent nanotube production was shown to be conserved across different species, signifying nanotubes as ubiquitous bacterial organelles ^17,20-22^. Electron microscopy (EM) and cryo-electron tomography (cryo-ET) ^23^ unveiled *B. subtilis* nanotubes to comprise chains of membranous segments with a continuous lumen, directly stemming from the cell membrane and traversing the cell wall ^8^. Nanotubes can be viewed as “intercellular” or “extending”, with the latter projecting from the producing cell into the surrounding environment ^8,18^. Notably, nanotube-like structures were shown to be produced by numerous Gram-positive and Gram-negative bacterial species [e.g., ^12,24-30^]. Moreover, our previous findings demonstrated the existence of bacterial outer membrane nanotube-like projections in thirteen different Gram-negative species ^31^. These observations highlight the potential of nanotubes to broadly impact bacterial physiology, communication, and adaptation across diverse ecological niches.

Here, we sought to elucidate how bacterial nanotubes are generated and how they establish intercellular connections. By integrating a tailored live-cell imaging strategy with *in situ* cryo-electron tomography, we directly visualized nanotube biogenesis and dynamics at high spatial and temporal resolution. This approach enabled us to capture the full sequence of nanotube projection, recipient-surface scanning, and docking to form an intercellular bridge. Furthermore, we uncovered the existence of a previously unrecognized bacterial membranous organelle, which we term tubeosome, that functions as a factory for nanotube biogenesis from which nanotubes emerge. Together, these findings provide a mechanistic framework for understanding how bacteria assemble membrane bridges for intercellular communication.

## Results

### Exposing nanotube networks in live bacteria

Nanotubes are difficult to visualize in living cells due to their thin diameter, fragility, and the absence of specific molecular markers. While nanotubes can be stained with a fluorescent membrane dye [e.g., ^8,12^], the intense signal emitting from the cell membrane largely obscures their visualization. To overcome this limitation, we devised a fluorescence microscopy methodology that focally separates cells and nanotubes, allowing clear nanotube monitoring using a fluorescent membrane dye (Figure 1A-1C; Figure S1A). *B. subtilis* cells, ectopically expressing YmdB to enhance nanotube production ^8,18^, were grown on a solid surface, a condition that promotes nanotube formation ^6,7^, supplemented with FM4-64 membrane stain. Fluorescence imaging exposed the existence of nanotube networks arrayed beneath the cells, with nanotubes spanning distances of several microns (∼1-15 µm), interconnecting proximal and remote cells, and located alongside cell chains (Figure 1D). The frequency of nanotube emergence varied from∼4-15% of cells (Figure S1C), likely an underestimate as only a subpopulation of focally separated nanotubes could be visualized. Often, nanotubes were associated with a pronounced fluorescent signal, indicating lipid-enriched sites (Figure 1D). Importantly, nanotube structures were nearly absent from a strain lacking the flagellar CORE components (Figure S1B-S1C), required for nanotube biogenesis ^17,20^. Phase-contrast images indicated that nanotube-producing cells appeared healthy and intact (Figure 1B, 1D). Nevertheless, to further validate their cellular integrity, we followed nanotubes formed by bacteria expressing cytoplasmic GFP, wherein the GFP signal reports cellular integrity and viability. Time-lapse microscopy affirmed that nanotube-producing cells were unharmed and continued to grow and divide (Figure S2A).

**Figure 1.**
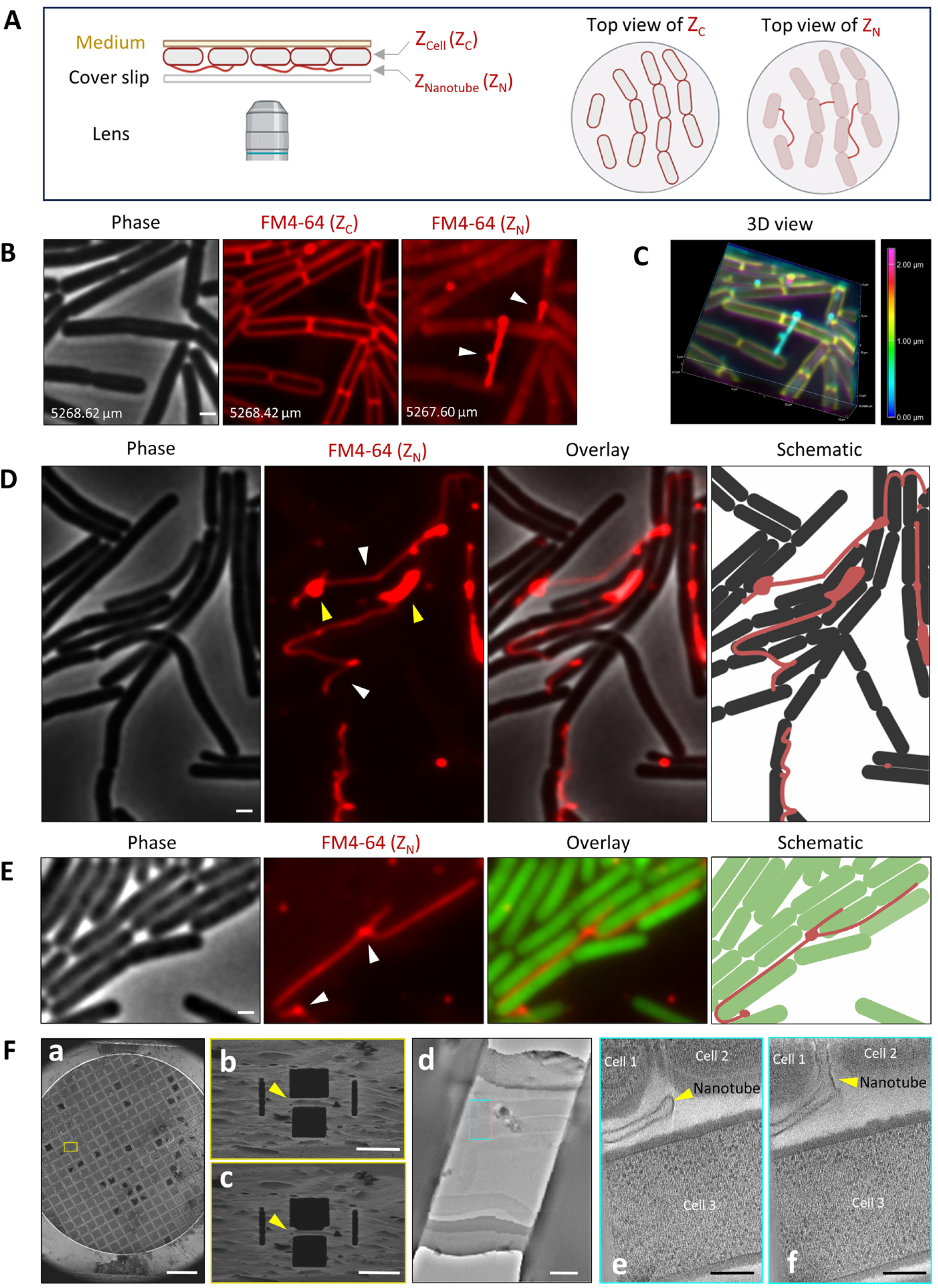
Revealing nanotube complexity in living bacteria. **(A)** A schematic representation of the designed methodology portraying the assembled apparatus employed to separate cell outlines and nanotubes following fluorescence membrane staining. Visualization was devised to focally separate cell membrane (Z_Cell,_ Z_C_) and nanotubes (Z_Nanotube,_ Z_N_), with top view of Z_C_ and Z_N_ depicted. Scale bar, 0.5 μm. **(B)** GB168 (P_IPTG_*-ymdB*) cells were grown on a solid LB medium containing FM4-64 lipophilic membrane dye that stains both nanotubes and cell membrane. After 60 min of incubation, Z stacks (0.2 µm steps) were collected to separate Z_C_ from Z_N._ Shown are phase contrast (gray), and FM4-64 (red) in Z_C_ and Z_N_ fluorescence images. Arrowheads highlight nanotubes. Z positions (µm) are indicated. Scale bar, 0.5 μm. **(C)** 3D deconvolution of Z-stack images shown in Fig. S1A. Deconvolution was performed using NIS-Element software (Nikon). Colour bar represents the depth of Z stacks at different focal planes. Cyan colour represents nanotubes. **(D)** GB168 (P_IPTG_*-ymdB*) cells were grown as in (B). Shown are phase contrast (gray), FM4-64 (red) in Z_N_ fluorescence, their overlay, captured after 60 min of incubation. Arrowheads highlight nanotubes (white), and lipid-enriched sites (yellow). Right panel depicts schematic of cells (black) and nanotubes (red). A representative field out of 3 independent biological repeats. Scale bar, 0.5 μm. **(E)** MG200 (P_IPTG_*-ymdB,* P1*_rrnB_-gfp*) cells, expressing GFP, were grown as in (B). Shown are phase contrast (gray), FM4-64 (red) in Z_N_ fluorescence, an overlay of FM4-64 (Z_N_) with GFP, captured after 60 min of incubation. Arrowhead highlights nanotube junctions. Right panels depict schematics of cells (green) and nanotubes (red). Representative fields out of 3 independent biological repeats. Scale bar, 0.5 μm. **(F)** MG155 (Δ*ponA*, P_IPTG_*-ymdB*, Δ*hag*) were grown on gold electron microscopy grids on a solid LB medium and then plunge-frozen with subsequent cryo-FIB milling and cryo-ET imaging. (a) Grid overview in SEM, before cryo-FIB-milling. (**b**, **c**) gradual FIB-milling of cell clusters to produce lamellae amenable for cryo-ET imaging (yellow arrows). (**d**) 2D cryo-EM image of the FIB-milled lamella before data acquisition, blue rectangle marks the area presented in images e and f. (**e, f**) Slices at different Z-heights through a cryo-ET of a lamella showing three bacterial cells with a nanotube, marked with the yellow arrowhead, extending between them. Scale bars, (a) 500 μm, (b-c) 10 μm, (d) 1 μm, (e-f) 200 nm.

A deeper investigation of nanotube patterns exposed prominent long membranous structures, often aligned with the typical chains of *B. subtilis* cells, as if they were embedded routes within the bacterial community (Figure 1E). Frequently, junctions were formed between nanotubes at cell intersections, with some nanotube clusters entangled together, denoting complex intercellular networks (Figure 1E; Figure S2B). In line with our earlier EM findings ^8^, two distinct types of nanotubes were evident: intercellular bridging nanotubes, tethered at both ends, connecting neighbouring cells (Figure 1B-1E), and extending nanotubes, originating from one cell and featuring a free disengaged end (Figure S2C). Notably, lipid-enriched sites were mostly associated with bridging nanotube ends, extending nanotube emanation sites and free tips, as well as with nanotube junctions (Figure 1D-1E; Figure S2B-S2C). Similar nanotube morphologies were formed in an interspecies manner between *B. subtilis* and its close relative *B. megaterium* (Figure S2D-S2E), signifying the potential utilization of these structures as major interspecies communication paths.

To further substantiate our fluorescence microscopy findings, we visualized nanotubes extending between cells using cryo-focused ion beam (cryo-FIB) milling of *B. subtilis* cell clusters grown on a solid surface. Cryo-FIB milling enables thinning of otherwise inaccessible thick samples to generate lamellae suitable for high-resolution cryo-ET data collection ^23^. Cryo-tomograms revealed membranous structures positioned between cells and extending across multiple z-planes of the reconstructed volumes (Figure 1F). These structures exhibited morphologies consistent with those observed by live-cell imaging, providing independent structural support for the fluorescence-based observations (Figure 1F).

### Real-time dynamics of intercellular and extending nanotubes

We next tracked real-time dynamics of nanotubes at millisecond intervals over extended periods (minutes) through time-lapse microscopy. Networks of nanotubes were rapidly moving, indicating their remarkable plasticity (Figure 2A; Movie S1). Detailed observation of intercellular nanotube dynamics revealed that the cell-anchored regions, referred to as "knots" (Figure 2B-2C, green), remained relatively stationary. Though, the elongated tubular regions (Figure 2B-2C, cyan, yellow) exhibited wave-like undulating motion on a timescale of seconds, with an amplitude ranging from approximately 0.5 to 1.8 µm (Figure 2B-2C; Movie S2). Furthermore, time-lapse microscopy of extending nanotubes uncovered their highly spatial dynamics in multiple directions, occasionally approaching neighbouring cells (Figure 2D; Movie S3). Following the trajectories of an extending nanotube free tip (Figure 2E, blue circle) compared to knot positions (Figure 2E, magenta and green circles), depicted a rapid movement of the free tip in both X and Y directions, with velocity reaching up to 0.5 µm/s, while knot positions were almost stationary (Figure 2E-2F). We conclude that bridging nanotubes are anchored at their tips and knots, while exhibiting undulating motion. Furthermore, extending nanotubes demonstrate elasticity and rapid multidirectional movement.

**Figure 2.**
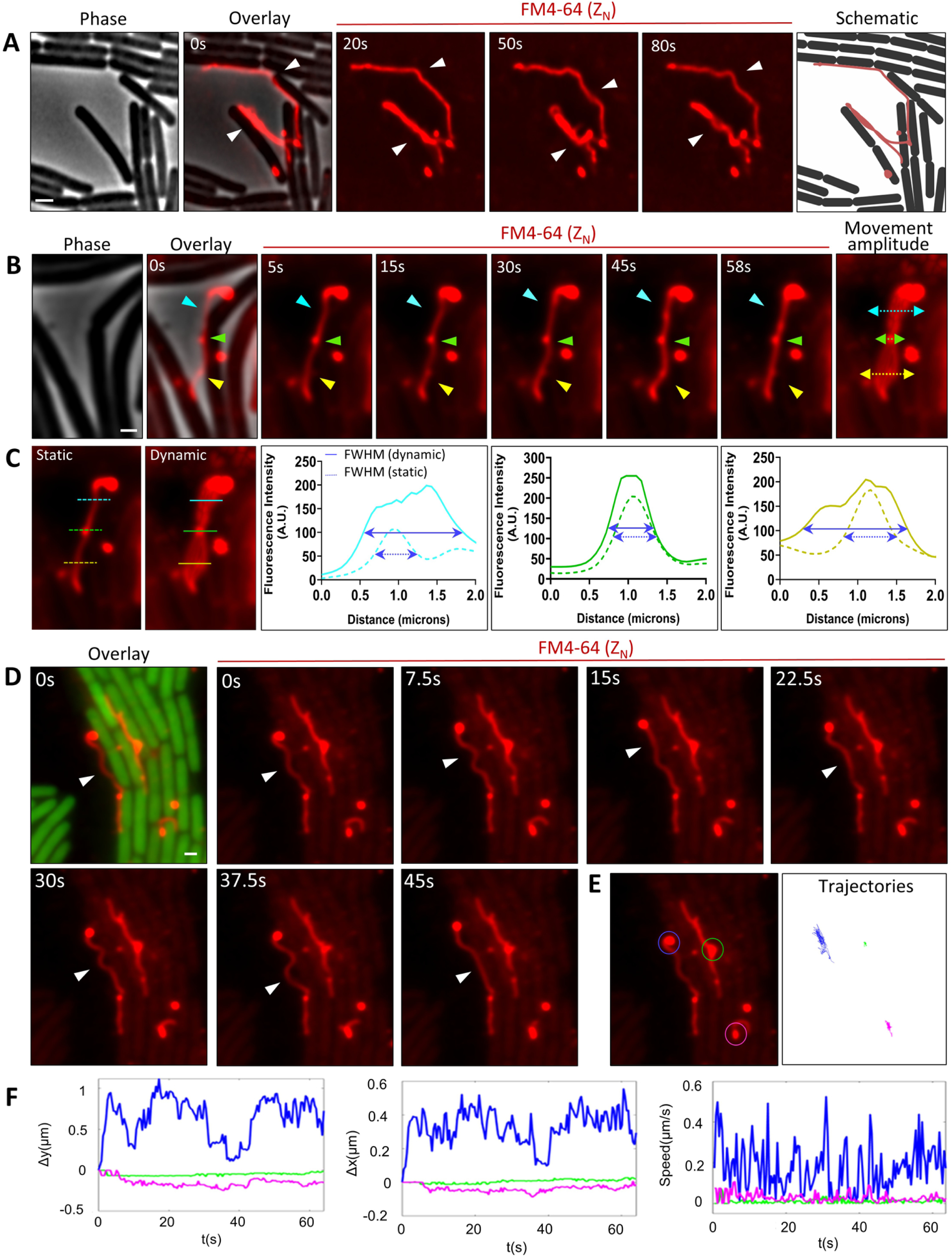
Intercellular and extending nanotube dynamics. **(A)** GB168 (P_IPTG_*-ymdB*) cells were grown on a solid LB medium containing FM4-64. After 90 min of incubation, fluorescence from FM4-64 (red) in Z_N_ was recorded (300 ms/frame, total 83 s). Shown are phase contrast (gray), an overlay of phase with FM4-64 (red) in Z_N_ (t=0), FM4-64 in Z_N_ images capturing the motion of nanotube dynamics at the indicated time points, and schematic depicting cells (black) and nanotubes (red). See corresponding Movie S1. Scale bar, 1 μm. **(B)** GB168 (P_IPTG_*-ymdB*) cells were grown as in (A). After 90 min of incubation, fluorescence from FM4-64 (red) in Z_N_ was recorded (200 ms/frame, total 58 s). Shown are phase contrast (gray), an overlay of phase with FM4-64 (red) in Z_N_ (t=0), FM4-64 in Z_N_ images capturing the motion of intercellular nanotube dynamics at the indicated time points, and the nanotube movement amplitude determined by ImageJ. Arrows highlight elongated tubular regions (cyan, yellow) and the "knot" region (green). See corresponding Movie S2. Scale bar, 0.5 μm. **(C)** Shown are nanotube movement amplitudes in static and dynamic states. Dashed (static state) and solid (dynamic state) lines highlight elongated tubular (cyan, yellow), and "knot" (green) regions. The fluorescence intensity profiles of static and dynamic nanotube movements are presented. Corresponding graphs show the fluorescence intensity (Y axis, A.U.) as a function of distance from 0-2 µm (X axis). Dashed blue lines in graphs indicate the full width at half maximum (FWHM) for static movement in a single frame, and the solid blue lines represent the FWHM average of sub-images for dynamic movements. See corresponding Movie S2. **(D)** MG200 (P_IPTG_*-ymdB,* P1*_rrnB_-gfp*) cells, expressing GFP, were grown as in (A). After 90 min of incubation, FM4-64 (red) in Z_N_ was recorded at (300 ms/frame, total 64 s). Shown are an overlay of GFP (green) with FM4-64 (red) in Z_N_ (t=0), and FM4-64 in Z_N_ images capturing the motion of nanotube dynamics at the indicated time points. Arrowhead highlights extending nanotube dynamics. See corresponding Movie S3. Scale bar, 0.5 μm. **(E)** Left panel [(D), FM4-64 in Z_N_ (t=0)] shows extending nanotube free tip (blue circle), and nanotube connected tips (magenta and green circles). Right panel shows trajectories of the corresponding nanotube regions. ImageJ application was used to track and plot the circled regions in each frame for the course of 60 s. See corresponding Movie S3. **(F)** Shown are position-time graphs of Y (left) and X (middle) directions, along with a graph of the absolute velocity (speed, µm/s) (right) over time (s), corresponding to the circled regions in (E).

### Nanotube docking on a recipient bacterium: the formation of an intercellular bridge

Thus far, real-time establishment of an intercellular nanotube bridge has not been captured, prompting the utilization of our live imaging system to monitor the occurrence of this critical event. We reasoned that extending nanotubes might represent an intermediate step toward the formation of an intercellular bridge and therefore focused on tracking their motion. Monitoring extending nanotube dynamics unveiled their engagement in a prolonged process, lasting several minutes, during which their free tips were repeatedly touching, brushing against, and stroking the neighbouring cell surface (Figure 3A; Figure S3A; Movie S4). Following numerous attempts, a successful intercellular connection was established, as indicated by an immediate switch in nanotube tip dynamics from rapid to constrained secured knot (Figure 3B-3C; Movie S4). In some cases, we could capture nearly successful attempts by a nanotube producer to establish a connection with its targeted cell. Throughout these events, connections appeared to persist for a few seconds but ultimately collapsed (Figure S3B). Interestingly, the establishment of an intercellular connection could coincide with a significant shortening of the nanotube length, accompanied by a pronounced increase in membrane fluorescence signal along the nanotube (Figure 3D-3E; Movie S5). This observation highlights the high degree of elasticity and flexibility inherent in these membranous structures.

**Figure 3.**
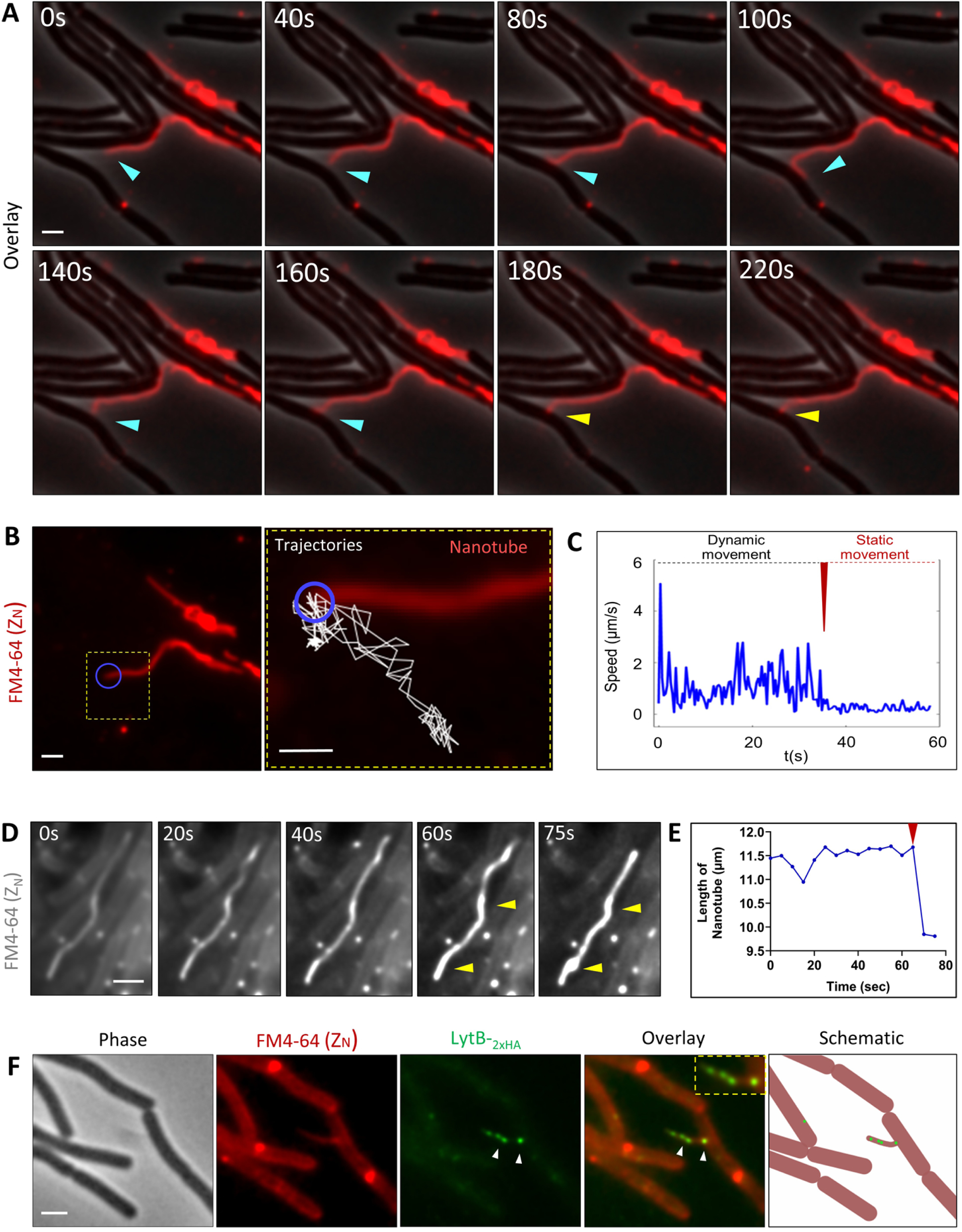
Visualizing the establishment of an intercellular nanotube bridge. **(A)** GB168 (P_IPTG_*-ymdB*) cells were grown on a solid LB medium containing FM4-64 membrane dye. After 100 min of incubation, fluorescence from FM4-64 (red) in Z_N_ was recorded at (200 ms/frame, total 238 s). Shown are overlays of phase contrast (gray) with FM4-64 (red) in Z_N_, captured at the indicated time points. Arrowheads highlight nanotube dynamics (cyan) and penetration site (yellow). A representative experiment out of 4 similar independently captured events. See corresponding Movie S4. **(B)** Shown are FM4-64 (red) in Z_N_ image [(A), FM4-64 in Z_N_ (t=0)] (left panel), and enlargement of the inset displaying nanotube trajectory analysis (right panel). Movement trajectories (white) of the nanotube tip (blue circle), ranging from 1-215 frames were analyzed. See corresponding Movie S4. **(C)** Shown is the absolute velocity (speed, µm/s) of nanotube tip trajectories, corresponding to the blue circle in (B). Red arrowhead highlights an intercellular bridging event. See corresponding Movie S4. **(D)** GB168 (P_IPTG_*-ymdB*) cells were grown as in (A). After 100 min of incubation, fluorescence from FM4-64 in Z_N_ was recorded (200 ms/frame, total 75 s). Shown are FM4-64 in Z_N_ images captured at the indicated time points. Arrowheads highlight increasing fluorescence signal sites along nanotubes over time. See corresponding Movie S5. **(E)** A graph representing the length of the nanotube in (D) over time, measured using ImageJ. Red arrowhead highlights an intercellular bridging event associated with nanotube shortening. See corresponding Movie S5. **(F)** AB266 (*lytB-_2XHA_*) cells were grown as in (A). Shown are images of phase contrast (gray), FM4-64 (red) in Z_N_, anti-HA Qdot525 labeled LytB molecules (green), an overlay of FM4-64 in (Z_N_) with anti-HA Qdot525 labeled LytB molecules, captured after 60 min of incubation. Arrowheads highlight localization of LytB molecules along an extending nanotube. Inset image (Overlay) presents the enlargement of this nanotube decorated with LytB molecules. Right panel depicts schematic of cells and an extending nanotube(red), with arrayed LytB molecules (green). Scale bars, 1 μm.

The relatively lengthy process undertaken by extending nanotubes to establish a stable intercellular connection raised the hypothesis that, during this phase, extending nanotubes deliver lytic enzymes onto the surface of neighbouring bacteria to locally compromise their cell wall. One such enzyme could be LytB, implicated in puncturing the recipient cell wall to facilitate nanotube penetration ^19^. Thus, we visualized surface-located single LytB molecules utilizing quantum dot (QD)-labeled antibodies (Figure S4A). Employing live immunofluorescence, we could indeed detect intercellular transfer of LytB molecules in a CORE-dependent manner (Figure S4B-S4D), consistent with our previous EM observations ^19^. Moreover, LytB molecules were found to be arrayed along extending nanotubes (Figure 3F), substantiating that during their motion, nanotubes deposit lytic proteins over the recipient cell wall.

### Nanotubes emerge from lipid accumulation sites

To characterize nanotube emanation sites, we took advantage of the findings that the flagellar CORE complex serves as a platform for nanotube biogenesis ^20^. Hence, the primary CORE component, FlhA, encompassing a cytoplasmic ectodomain, was fused with Stay-Gold (SG) fluorescent protein ^32^, serving as the sole chromosomal *flhA* copy. The resultant fusion protein was functional, as signified by the retained motility of the cells (Figure S5A). FlhA-SG was found to localize into distinct bright foci on the bacterial membrane, marking CORE complexes (Figure S5B). Co-visualizing FlhA-SG and nanotubes unveiled detectable FlhA-SG foci positioned at one end of bridging nanotubes (Figure 4A), likely denoting sites of nanotube emergence, reinforcing our current understanding ^20^. Notably, FlhA-SG foci frequently colocalized with lipid-enriched sites, as evidenced by heightened fluorescence signal of the membrane dye (Figure 4A). In some instances, FlhA-SG foci and lipid accumulation sites were associated with a prominent nanotube, while in others, no apparent nanotube was present (Figure 4A), inferring at the potential function of these sites as reservoirs for future nanotube projection. To further investigate this possibility, we sought at capturing nanotube projection events in real time. Remarkably, we could successfully record the rapid firing of nanotubes, extending to a distance of several microns within few msec (Figure 4B; Movie S6). Nanotubes originated from lipid-enriched sites, which significantly diminished following the protrusion event (Figure 4B; Movie S6), suggesting that these sites serve as lipid storage compartments that facilitate rapid nanotube extension.

**Figure 4.**
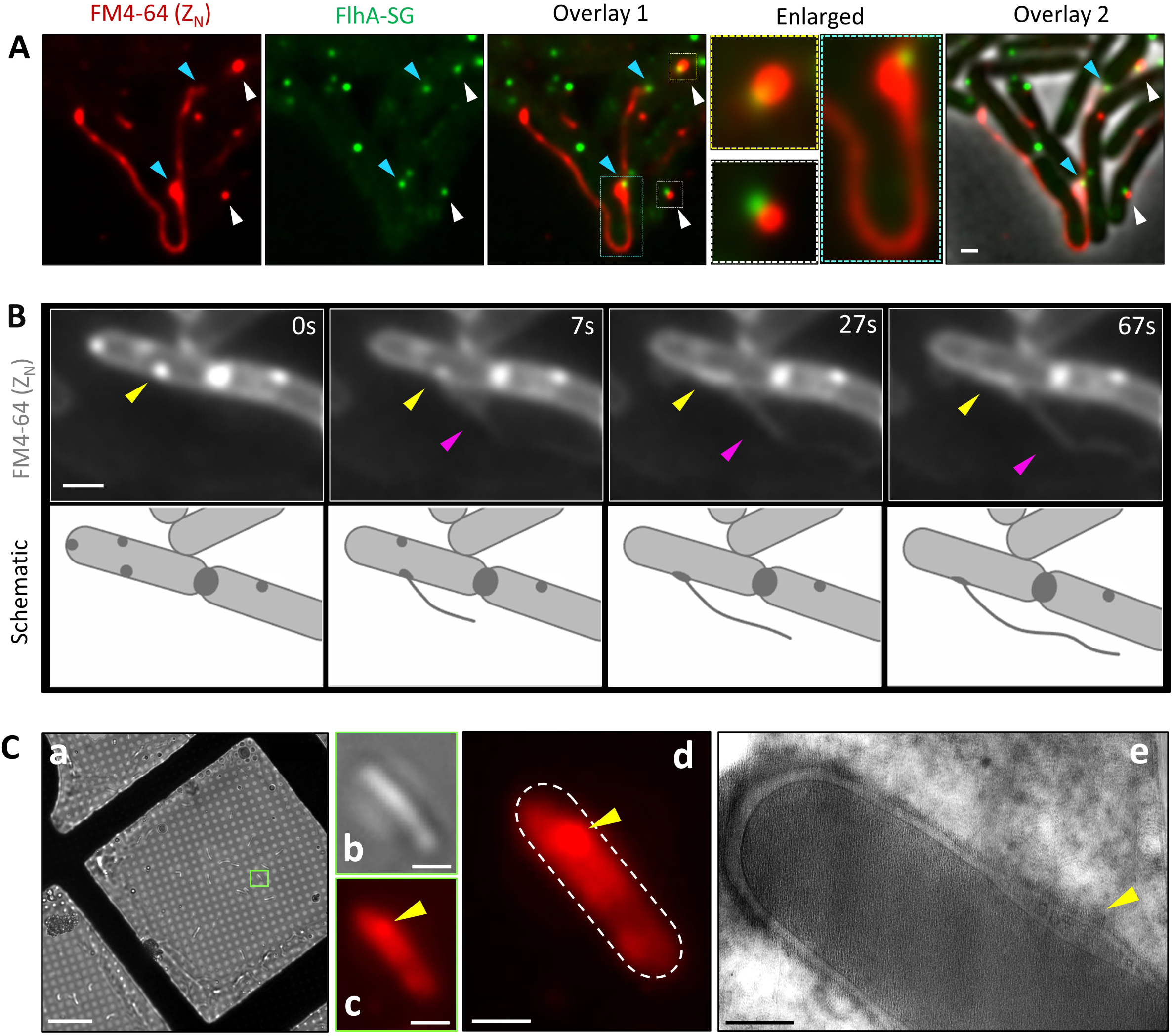
Nanotube emerge from sites of membrane accumulation. **(A)** GV594 (*flhA-lin_4x_-sg,* P_IPTG_*-ymdB*) cells were grown on a solid LB medium containing FM4-64. Shown are fluorescence images of FM4-64 (red) in Z_N_, FlhA-SG (green), overlay 1 of FM4-64 in Z_N_ with FlhA-SG, overlay 1 inset enlargements, and overlay 2 of FM4-64 in Z_N_, FlhA-SG, and phase contrast (gray), captured after 60 min of incubation. Blue arrowheads highlight nanotube emanation sites, and white arrowheads highlight membrane accumulation sites, both colocalize with FlhA-SG foci. Scale bar, 0.5 μm. **(B)** GB168 (P_IPTG_*-ymdB*) cells were grown as in (A). After 100 min of incubation, fluorescence from FM4-64 in Z_N_ was recorded (400 ms/frame, total 235 s). Upper panels show FM4-64 in Z_N_ images captured right before and after nanotube protrusion event, at the indicated time points. Yellow arrowheads highlight nanotube protrusion sites, and pink arrowheads highlight emerging nanotubes. Bottom panels depict schematics of cells and the emerging nanotube. A representative experiment out of 3 similar independently captured events. See corresponding Movie S6. Scale bars, 0.5 μm. **(C)** MG155 (Δ*ponA*, P_IPTG_*-ymdB*, Δ*hag*) were grown on electron microscopy grids on a solid LB medium containing FM4-64 and then plunge-frozen and imaged with cryo-light microscopy. Panel (**a**) is a bright field cryo-light microscopy of frozen cells on the grid. Panels (**b, c**) are bright-field and fluorescent images, respectively, of the cell highlighted by the green box in panel (a). Panel (**d**) is an enlarged fluorescent image of the cell shown in panels (b) and (c). Panel (**e**) represents a slice through a cryo-electron tomogram of the cell shown in panels (b-d). Yellow arrowheads in panels (c), (d), and (e) highlight the fluorescent focus in the cell and the corresponding location in the cryo-electron tomogram where a tubeosome can be seen. scale bars, (a) 20 μm, (b-d) 0.5 μm, (c) 200 nm.

### The existence of a tubeosome organelle

To assess the structure of the membrane-enriched nanotube emanation sites at higher resolution, we carried out *in situ* cryo-ET analysis. Given the relatively thick nature of *B. subtilis* cells, we employed a Δ*ponA* mutant, which is thinner than WT, amenable for cryo-ET imaging ^33-36^. Correlative Light and Electron Microscopy (CLEM) exposed that the membrane-enriched sites correspond to previously uncharacterized membrane invagination and vesicular densities (Figure 4C). Examination of the cell population showed that these membrane structures varied from small invaginations to elaborate multilayered rolled membranes, exhibiting X- and Y-axis dimensions consistent with an approximately spherical shape and diameters ranging from 25 to 400 nm (Figure 5A and 5F; Figure S6A-S6C). This size variation may correspond to different stages in the assembly of these structures. Such membranous layers were present in 43.2% of the surveyed cells (Figure 5G), with some containing more than a single structure (Figure S7A). Consistently, similar membranous structures were clearly seen in ultrathin sections using conventional Transmission Electron Microscopy (TEM) (Figure S7B). As these membranous assemblies resemble fire hose-like rolled nanotubes, we termed these organelles “tubeosomes”. 3D segmentations showed that tubeosomes formed in an atypical location, a pseudo-periplasmic-like compartment between the plasma membrane and the cell wall (Figure 5B; Figure S6A-S6C; Movie S7). At the resolution of our cryo-tomograms, multiple tubeosome compartments seemed to be arranged side-by-side in an orderly fashion beneath the cell wall (Figure 5C-5E; Movie S8). As nanotubes extend from the cell within milliseconds (Figure 4B), capturing their emergence from tubeosomes by cryo-ET is unlikely. Nevertheless, one of the folded tubeosomes exhibited nanotube-like extension (Figure 5E, red arrowhead; Movie S8), suggesting that tubeosomes serve as nanotube-manufacturing organelles.

**Figure 5.**
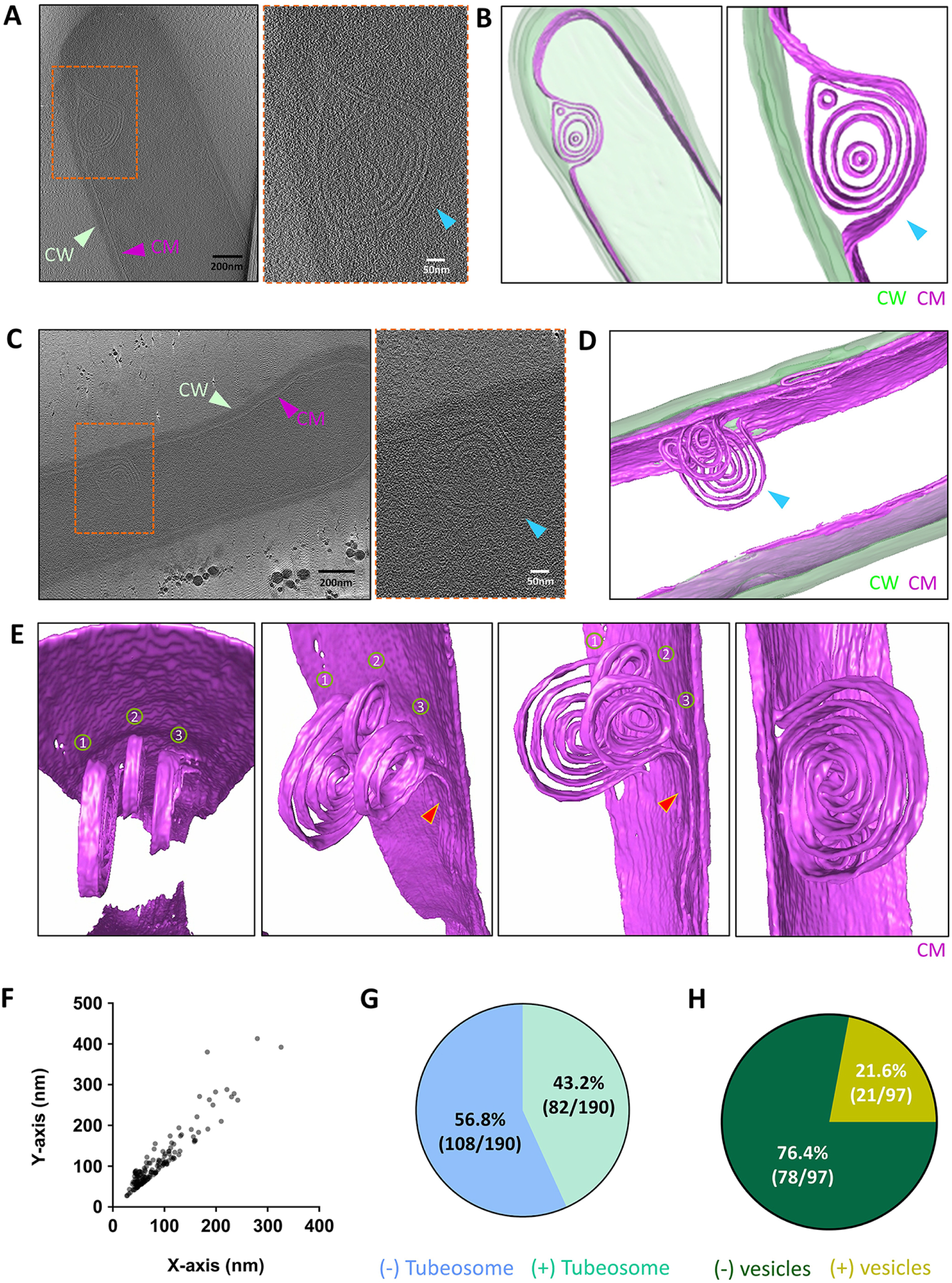
Characterization of tubeosome organelles. **(A)** AB328 (Δ*ponA*, P_IPTG_*-ymdB*, Δ*hag*) cells were grown on solid LB medium and subjected to Cryo-ET. Left panel shows a slice through a cryo-electron tomogram of a cell indicating the presence of the tubeosome structure (dashed orange rectangle), while the right panel shows inset enlargement. Scale bars, 200 nm (left panel), and 50 nm (right panel). See corresponding Movie S7. **(B)** 3D segmented volume of (A). Left panel shows an overview of the cell with the tubeosome, while the right panel shows zoomed-in views of tubeosome structure. See corresponding Movie S7. **(C)** Left panel shows a slice through a cryo-electron tomogram of a cell harboring stacks of membrane invaginations corresponding to the tubeosome structure, while the right panel shows inset enlargement. Scale bars, 200 nm (left panel), and 50 nm (right panel). See corresponding Movie S8. **(D)** 3D segmentations for the cell shown in (C) to highlight the tubeosome. See corresponding Movie S8. **(E)** 3D segmentations for the cell shown in (C), rotated at different angles, displaying enlarged images of the tubeosome region, highlighting 3 parallel manufactured tubeosomes (1,2,3) from different angles, with presumably a nanotube emerging from tubeosome (3). See corresponding Movie S8. **(F)** A size distribution of tubeosome dimensions in X and Y axis (nm) from cells grown and analyzed as in (A). Each dot represents an individual tubeosome positioned according to its size (*n*_cells_ = 190). **(G)** A pie chart displaying the quantitative distribution (%) of cells with or without tubeosome structures from 190 of AB328 cells grown and analyzed as in (A) (*n*_cells_ = 190). **(H)** MG164 (Δ*ponA*, Δ*CORE*) cells were grown on solid LB medium and subjected to Cryo-ET. Shown is a pie chart displaying the quantitative distribution (%) of cells with or without multiple vesicle structures in the pseudo-periplasmic space between the membrane and the cell wall (*n*_cells_ = 97). 3D Segmentations: CW (cell wall; green), CM (cell membrane; purple). Arrowheads highlight CW (cell wall; green), CM (cell membrane; purple), tubeosome (blue), a potential emerging nanotube-like structure (red).

### Tubeosome formation is perturbed in the absence of CORE

Cryo-ET visualization of extending nanotubes confirmed that, as previously observed, they are composed of membrane stacks and fused chains of vesicles (Figure 6A) ^8^, with some of them appearing to share a continuous lumen at the resolution of our cryo-tomogram (Figure 6B; Movie S9). As nanotubes emerge from sites of CORE complexes, we next asked whether tubeosomes can be formed in mutants lacking CORE. Typical tubeosomes were almost absent from Δ*CORE* mutant cells (1/97); surprisingly however, a subset of cells (21.6%) contained pseudo-periplasmic compartments resembling tubeosomes but filled with multiple rows of membrane vesicles aligned along the membrane exterior rather than the stacked membranes observed in wild-type cells (Figure 5H; Figure 6C-6D; Figure S7C; Movie S10). These structures were hardly detected in wild-type cells (6/214, 2.8%). Given that nanotubes seem to be composed of fused vesicle chains, we infer that these vesicle-rich compartments might represent an intermediate stage in tubeosome formation, with CORE required for completion of nanotube biogenesis.

**Figure 6.**
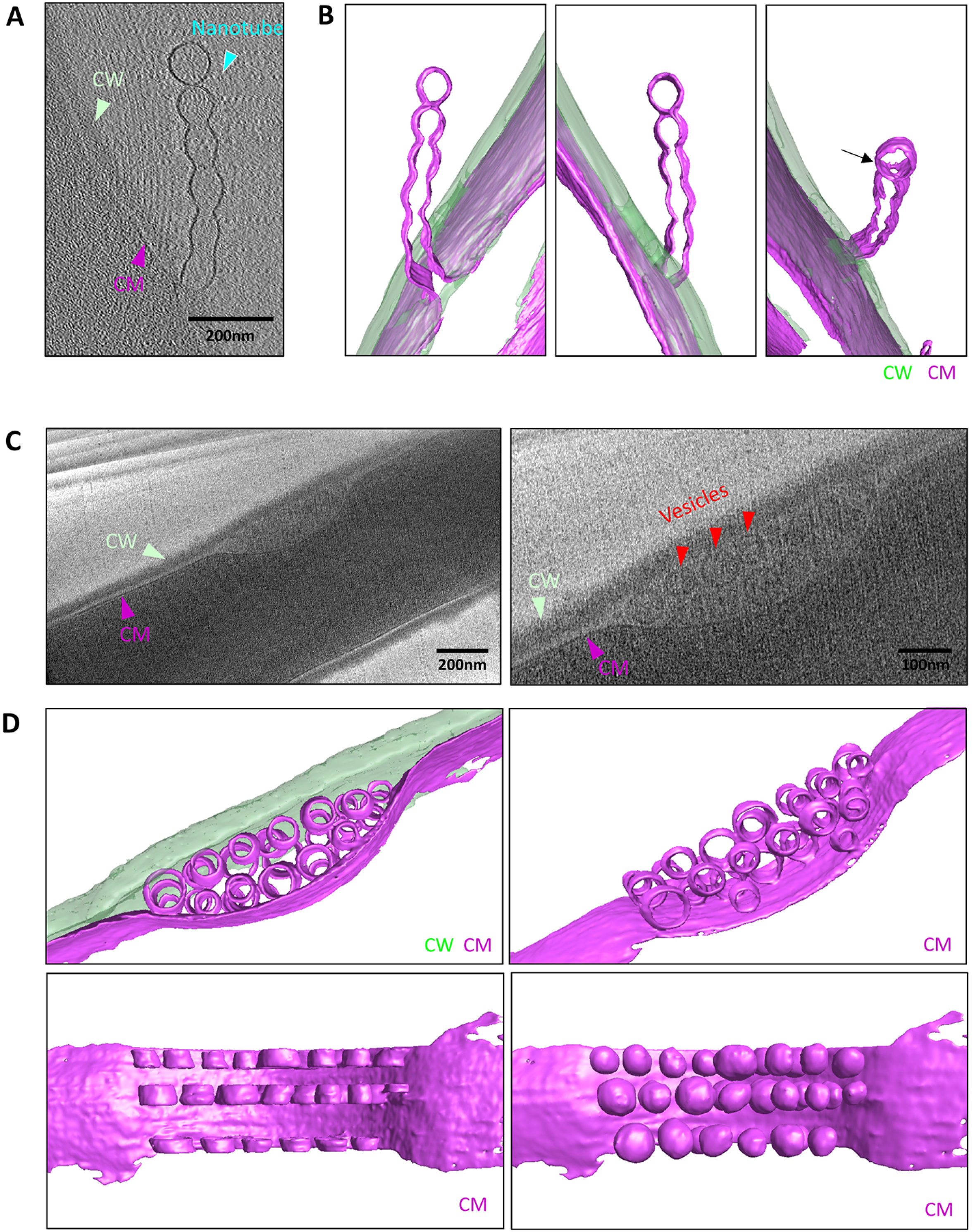
Δ*CORE* mutant cells lack tubeosomes but display stacks of membrane vesicles. **(A)** AB328 (Δ*ponA*, P_IPTG_*-ymdB*, Δ*hag*) cells were grown on solid LB medium and subjected to Cryo-ET. Shown is a slice through a cryo-electron tomogram displaying a cell harboring an extending membranous nanotube. See corresponding Movie S9. **(B)** 3D segmentations of the nanotube site in (A) rotated at different angels. An arrow highlights a hole that links the terminal vesicle segment with the nanotube fused vesicle chain, indicating a shared lumen. See corresponding Movie S9. **(C)** MG164 (Δ*ponA*, Δ*CORE*) cells were grown on solid LB medium and subjected to Cryo-ET. Left panel shows a slice through a cryo-electron tomogram of a cell harboring stacks of membrane vesicles, while the right panel shows the enlargement of the vesicle compartment. See corresponding Movie S10. **(D)** 3D segmentations of (C). Upper panels and lower left panel display clustered vesicles packed in a “sac-like” compartment underneath the cell wall rotated at different angles of the vesicle compartment. See corresponding Movie S10. Lower right panel shows a schematic interpretation of the 3D segmentation where the vesicles were manually completed into spheres for illustrative visualization only to compensate for the missing-wedge artifact. 3D Segmentations: CW (cell wall; green), CM (cell membrane; purple). Arrowhead highlights CW (cell wall; green), CM (cell membrane; purple), nanotube (cyan), and vesicles (red). Scale bars, 200 nm.

## Discussion

Despite their abundance, the process by which nanotubes are generated, projected, and bridge neighbouring bacteria has remained mysterious. Here, we unveil a previously undescribed bacterial organelle, the tubeosome, which serves for nanotube biogenesis. In the absence of CORE, tubeosome formation appears to be halted at an intermediate stage, as rows of membrane vesicles (Figure 7A). We thus propose that nanotubes may arise through the fusion of an array of membrane vesicles into elongated tubular structures, a process occurring within tubeosomes and is facilitated by the CORE complex (Figure 7A). Being a secretion apparatus, CORE may deliver proteins that catalyse this membrane-fusion process. Alternatively, the CORE could secrete factors necessary for nanotube stabilization and projection, such that in their absence, caged nanotubes collapse into vesicles. A few nanotubes can be produced within the same tubeosome site, suggesting a factory-like architecture for nanotube manufacture. Interestingly, the tubeosome resides in an unusual location, a curved membrane pocket situated in a pseudo-periplasmic space between the membrane and the cell wall, a domain that so far has not been defined as a bacterial cellular compartment. By employing a live-cell imaging strategy, we elucidated the cycle of nanotube biogenesis and dynamics, refuting a previous claim of nanotubes being a manifestation of cell death ^37^. Nanotubes rapidly extend from the tubeosomes, most likely via cell wall opening ^19^, and subsequently engage in a searching process, scanning and touching the surface of nearby bacteria for a period of a few minutes (Figure 7B). During this process, lytic enzymes, such as LytB, are deposited over potential recipient surfaces to locally pierce their cell wall (Figure 7B) ^19^. Once a breach is formed, the nanotube penetrates the membrane of the recipient bacteria, establishing a cell-to-cell membranous bridge. The formation of a secure intercellular connection is characterized by a sharp decrease in nanotube dynamics and velocity (Figure 7B).

**Figure 7.**
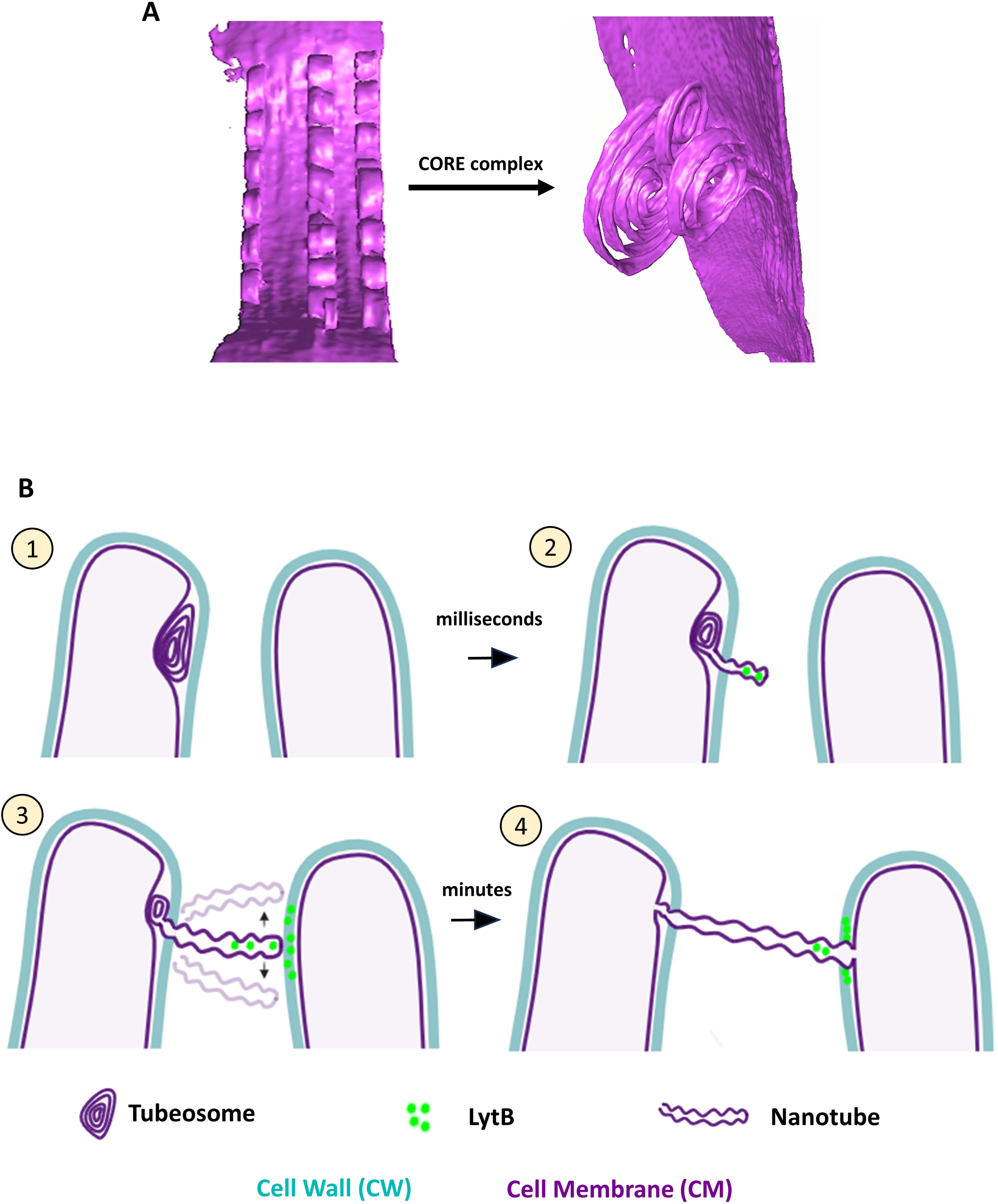
A model for nanotube formation, emergence, and intercellular bridging. **(A)** Based on our data, we proposed that tubeosome generation initiates with the formation of aligned membrane vesicles that, with the help of CORE, are converted into tubeosome-rolled tubes and are eventually released as nanotubes. Shown are 3D segmentations of vesicle chains (Δ*CORE,* left) and tubeosomes (WT, right). **(B)** A schematic model for the different stages of an intercellular nanotube formation: (1) the formation of nanotubes in a tubeosome organelle located in a pseudo-periplasmic space beneath the cell wall, (2) the rapid release of the nanotube within milliseconds from the tubeosome, (3) dynamic movement of the nanotube in a “scanning motion”, while depositing LytB molecules on the neighboring cell surface, and (4) the subsequent establishment of an intercellular nanotube bridge.

A few membranous organelles have been described in bacteria, including magnetosomes in *Magnetospirillum* species, thylakoid membranes in *Cyanobacteria*, and photosynthetic chromatophore in *Rhodobacter* ^38-44^. Additionally, cytoplasmic membranous stacks in *Coxiella burnetii* are thought to represent a membrane reservoir, accelerating the transition from a small to a large cell variant ^45,46^. Tubeosomes could be widespread bacterial organelles, as nanotubes have been observed across a broad spectrum of species [e.g., ^12,24-31^]. Curiously, tubeosomes morphologically resemble the previously described mesosome structures that were observed in both Gram-positive and -negative bacteria, but were eventually dismissed as artifacts resulting from EM fixation in the late 1970s [e.g., ^47-54^]. The occurrence of mesosomes, tubeosomes, and nanotubes in only a subpopulation, their transient nature and dependency on physiological conditions, may have contributed to inconsistencies in their detection, raising the possibility that tubeosomes and mesosomes may, in fact, represent related, or even the same, structures. It is conceivable that tubeosomes play broader roles in bacterial physiology, such as expanding the bacterial cell surface under stress conditions to support efficient membrane metabolism and nutrient acquisition. Notably, tubeosomes frequently localize near sites of septum formation, and thus may potentially serve as intracellular membrane reservoirs involved in septal membrane biogenesis. Tubeosomes or related structures could also account for the production of other membranous surface organelles, such as extracellular vesicle chains, known as nanopods, produced by *Caulobacter crescentus* and *Delftia* sp. Cs1–4 ^31,55^, or membranous nanowires involved in long-range extracellular electron transport generated by *Shewanella oneidensis* MR-1 ^56,57^. Furthermore, deliberate formation of membrane vesicles ^58,59^ may likewise be orchestrated through similar mechanisms, particularly since vesicle “sacs” were observed in cells lacking *CORE*.

Many questions regarding nanotube formation and dynamics remain to be elucidated. We have yet to determine how tubeosomes are assembled, and how they metamorphose to nanotubes, when nanotubes are released from tubeosomes, what is the source of energy fuelling their dynamics, and what cues do they recognize in recipient bacteria to establish a connection? Our study provides the first visualization of an intercellular bridging process in bacteria, revealing the presence of a tubeosome organelle and shedding light on nanotube biogenesis and dynamics. Given that nanotube-like structures have been reported to connect cells across diverse domains, including archaea and mammalian cells ^60-63^, the mechanism described here may represent a widely relevant strategy for intercellular communication.

## Acknowledgments

We thank the members of the Ben-Yehuda, Rosenshine, Kaplan, and Jensen laboratories for their insightful discussions. We are also grateful to Alex Rouvinski (Hebrew University) for helpful advice. VG was supported by the Planning and Budgeting Committee (PBC) of Israel postdoctoral fellowship, and SB by the Golda Meir postdoctoral fellowship. This work was funded by the European Research Council (ERC), Synergy grant (810186) awarded to SB-Y and IR, by the National Institute of General Medical Sciences (R35GM157116), and the Searle Scholars Program (SSP-2025-105) awarded to M.K, and by NIH grant RO1 (AI127401) awarded to GJJ.

## Author contributions

Conceptualization: MKG, TZ, GJJ, MK, IR, and SB-Y

Methodology: MKG, TZ, MZZ, AKB, RRP, VG, SB, OY, MR, DG, GJJ, IR, MK, and SB-Y

Investigation: MKG, TZ, MZZ, AKB, and RRP

Funding acquisition: GJJ, IR, MK, and SB-Y

Project administration: GJJ, IR, MK, and SB-Y

Supervision: GJJ, IR, MK, and SB-Y

Writing - original draft: MKG, TZ, GJJ, MK, IR, and SB-Y

## Supplemental Figure Legends

**Figure S1.**
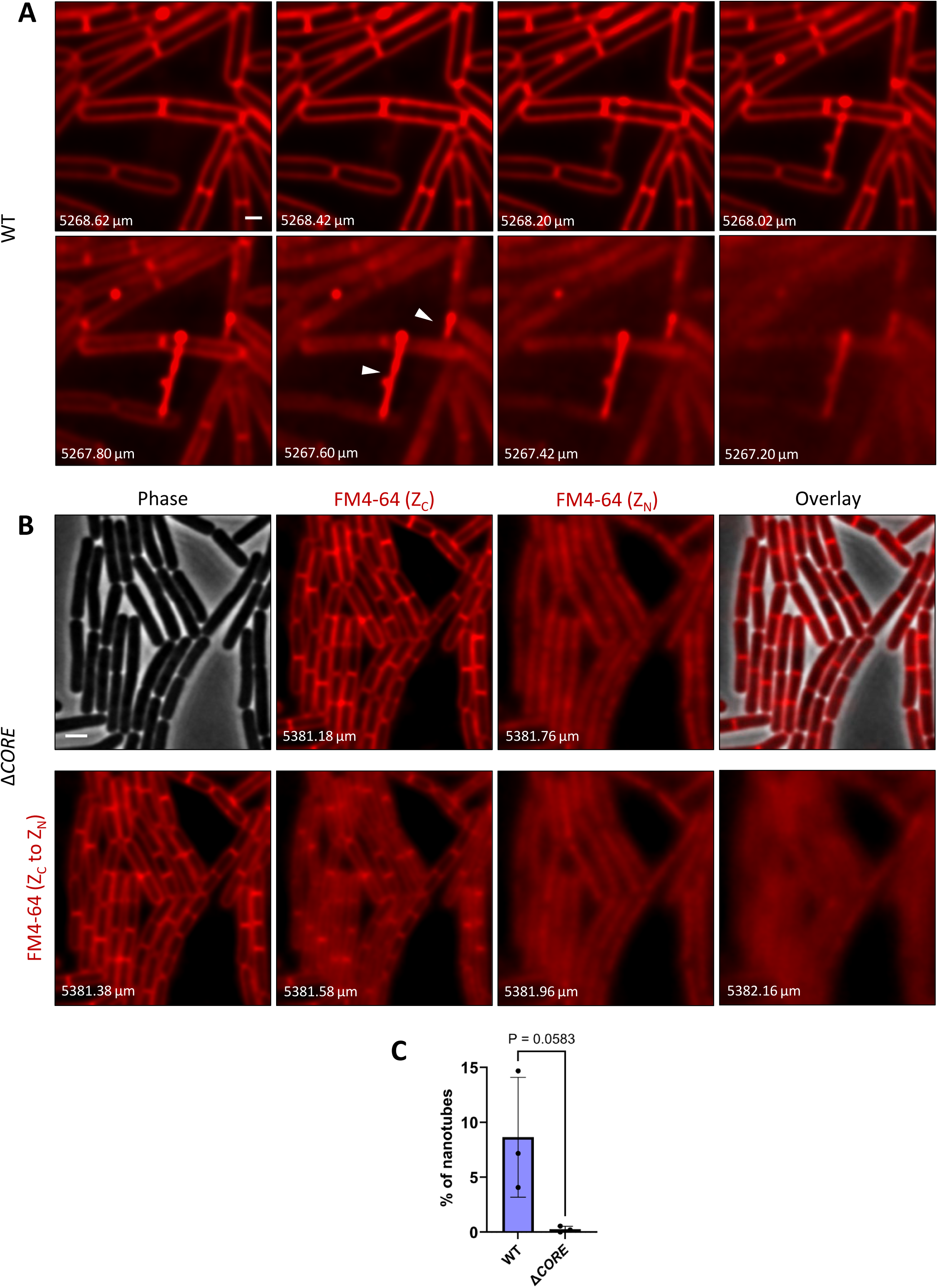
Nanotube visualization in specific focal planes. **(A)** GB168 (P_IPTG_*-ymdB*) cells were grown on a solid LB medium containing FM4-64, that stains both nanotubes and the cell membrane. After 60 min of incubation, Z stacks (0.2 µm) were collected to separate Z_C_ from Z_N._ Shown are FM4-64 (red) fluorescence images displaying the transition from Z_C_ to Z_N_. Arrowheads highlight nanotubes. Z positions (µm) are indicated. Scale bar, 0.5 μm. **(B)** SH9 (Δ*CORE*) cells were grown on a solid LB medium containing FM4-64. After 60 min of incubation, Z stacks (0.2 µm) were collected to separate Z_C_ from Z_N_. Upper panels show phase contrast (gray), FM4-64 (red) in Z_C_, FM4-64 in Z_N_, and overlay of FM4-64 in Z_C_ with phase. Bottom panels display FM4-64 fluorescence images through the transition from Z_C_ to Z_N_. Z positions (µm) are indicated. A representative field out of 3 independent biological repeats. Scale bar, 1 μm. **(C)** Quantitation of the experiments described in (A-B). Shown are the average number of nanotubes in WT (A) or Δ*CORE* (B). Nanotubes were manually counted using NIS analysis software. Each nanotube was counted as a single structure, although it usually spanned multiple cells. *P* values were determined using a one-tailed, unpaired Student’s *t*-test (*p* = 0.0583) Results were collected from 3 independent biological repeats for each strain, WT (*n*_cells_ =1244) and for Δ*CORE* (*n*_cells_ =1436).

**Figure S2.**
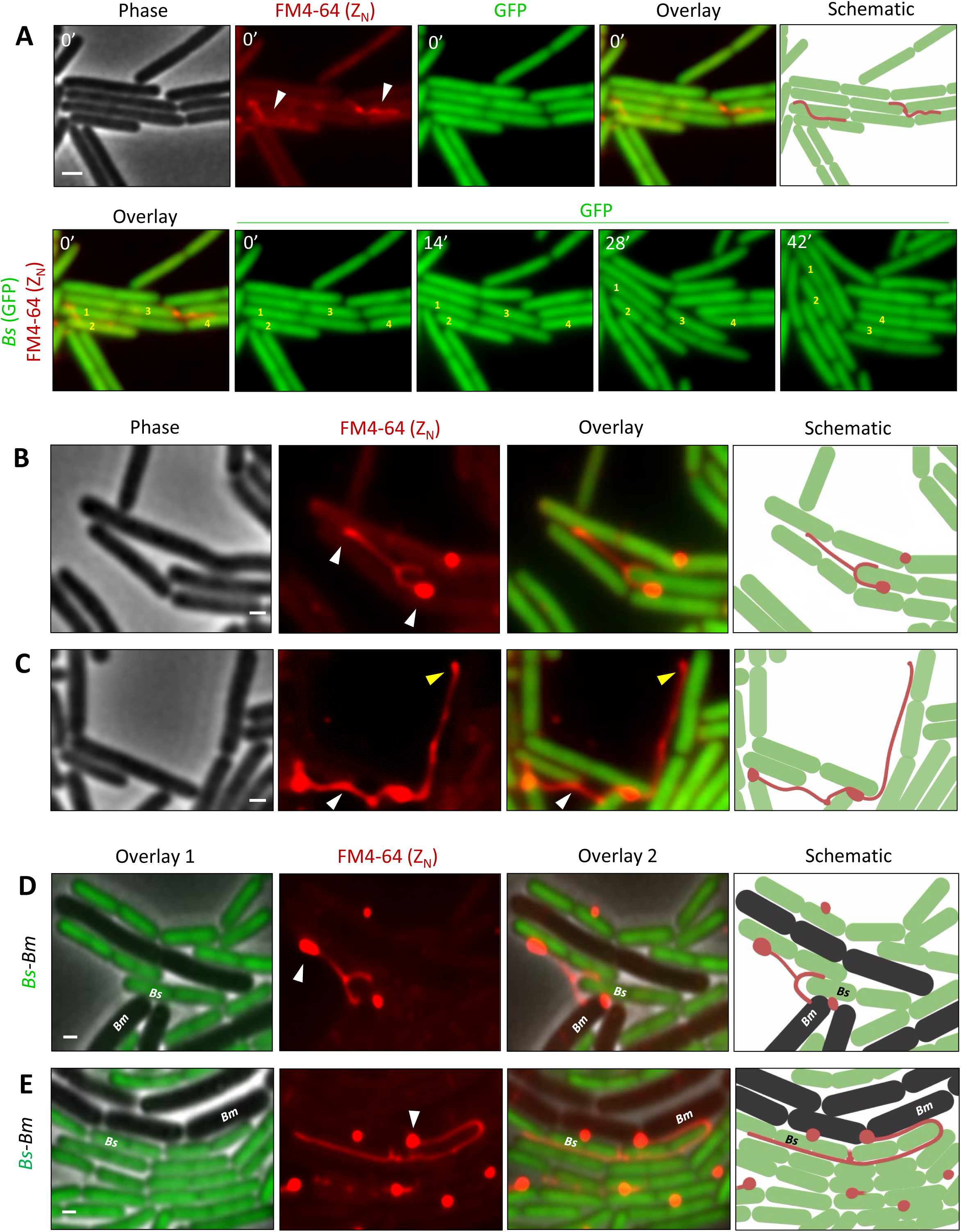
Nanotubes display both intra- and interspecies connections. **(A)** MG200 (P_IPTG_*-ymdB,* P1*_rrnB_-gfp*) cells were grown on a solid LB medium containing FM4-64. After 60 min of incubation, fluorescence from FM4-64 (red) in Z_N_ was captured (t=0), and fluorescence from GFP was followed at 7 min intervals for 60 min. Upper panels show phase contrast (gray), FM4-64 (red) in Z_N_, GFP (green), an overlay of FM4-64 in (Z_N_) with GFP, and a schematic depicting cells (green) and nanotubes (red). Bottom panels display an overlay of FM4-64 in Z_N_ with GFP at t=0 (overlay), and subsequent time-lapse images with fluorescence from GFP captured at the indicated time points (min). Cells labeled (1, 2) and (3, 4) represent pairs of nanotube-connected cells. Arrowheads highlight nanotubes. A representative experiment out of 2 similar independently captured events. **(B-C)** MG200 (P_IPTG_*-ymdB,* P1*_rrnB_-gfp*) cells were grown on a solid LB medium containing FM4-64 for 60 min. Shown are phase contrast (gray), fluorescence from FM4-64 (red) in Z_N_, an overlay of FM4-64 (Z_N_) with GFP. Arrowhead in **(B)** highlights lipid accumulation sites, whereas arrowheads in **(C)** highlight intercellular (white) and extending (yellow) nanotubes. Right panels depict schematics of cells (green) and nanotubes (red). Representative fields out of 3 independent biological repeats. **(D-E)** *B. subtilis* (*Bs*) MG57 (P_IPTG_*-ymdB,* P1*_rrnB_-gfp, ΔwapA*) cells (green) and *B. megaterium* (*Bm*) (WT) (gray) cells were mixed in 1:1 ratio, and grown on a solid LB medium containing FM4-64 for 60 min. *Bs* was deleted for *wapA* encoding a toxin, which is deleterious to *Bm*. Shown are overlay 1 of phase contrast (gray) with GFP (green), FM4-64 (red) in Z_N_, overlay 2 of phase contrast with GFP and FM4-64 in Z_N_, and schematics depicting cells and interspecies nanotubes. A representative experiment out of 3 independent biological repeats. Arrowheads highlight lipid accumulation sites. Scale bars, 0.5 μm.

**Figure S3.**
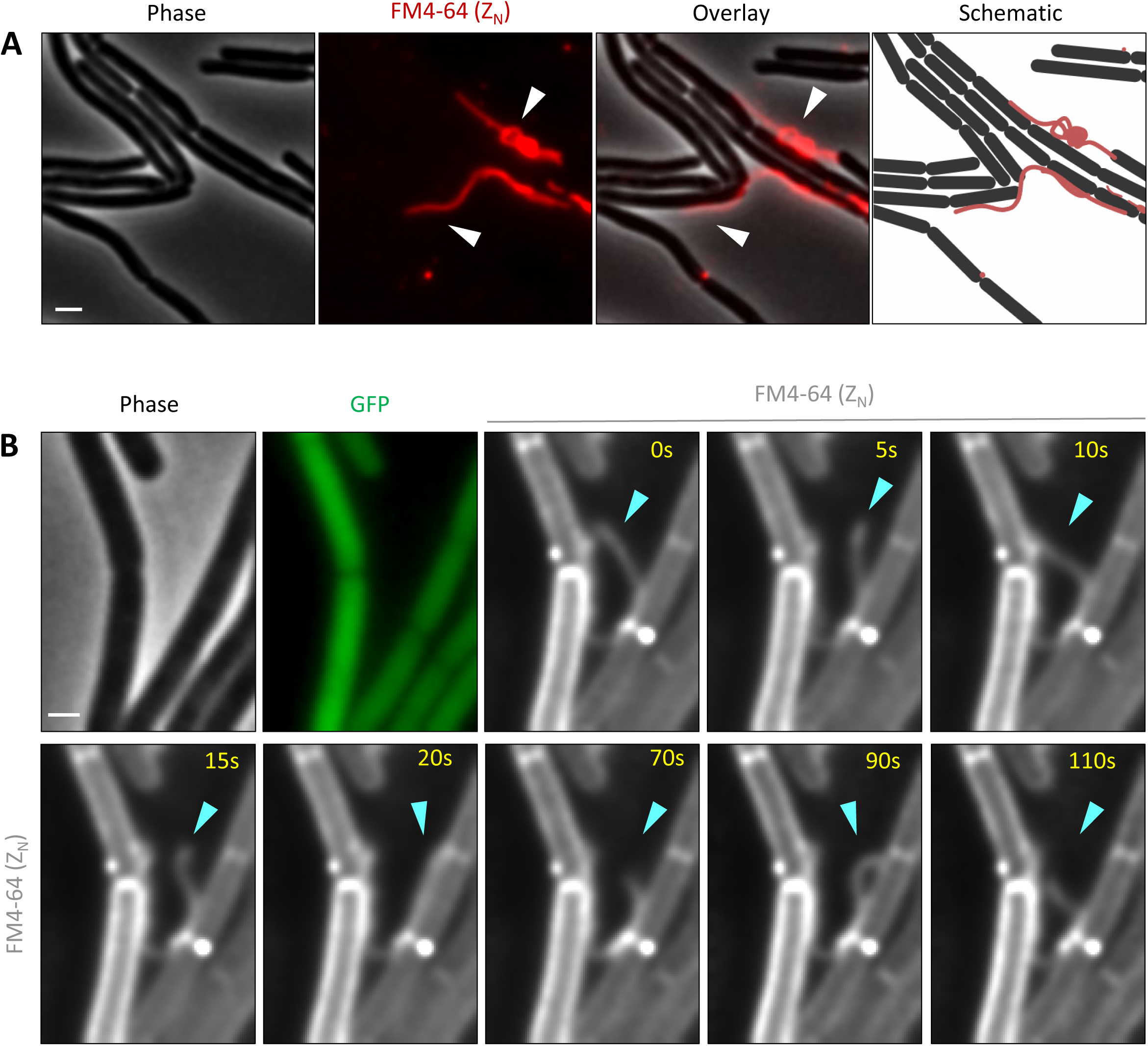
Dynamics of nanotube bridging events. **(A)** GB168 (P_IPTG_*-ymdB*) cells were grown on a solid LB medium containing FM4-64 membrane dye. Shown are phase contrast (gray), FM4-64 (red) in Z_N_ (t=0), overlay of FM4-64 in Z_N_ (t=0) with phase contrast, captured after 100 min of incubation, and schematic depicting cells and nanotubes. Arrowheads highlight nanotubes. Corresponding to Figure 3A and Movie S4. Scale bar, 1 μm. **(B)** MG200 (P_IPTG_*-ymdB,* P1*_rrnB_-gfp*) cells were grown on a solid LB medium containing FM4-64. After 100 min of incubation, fluorescence from FM4-64 in Z_N_ was recorded (200 ms/frame, 118 s) (capturing also Z_c_ in this case). Shown are phase contrast (gray), GFP (green), and a series of FM4-64 (white) in Z_N_, capturing the extending nanotube dynamics at the indicated time points. Arrowhead highlights the extending nanotube tip. Scale bar, 0.5 μm.

**Figure S4.**
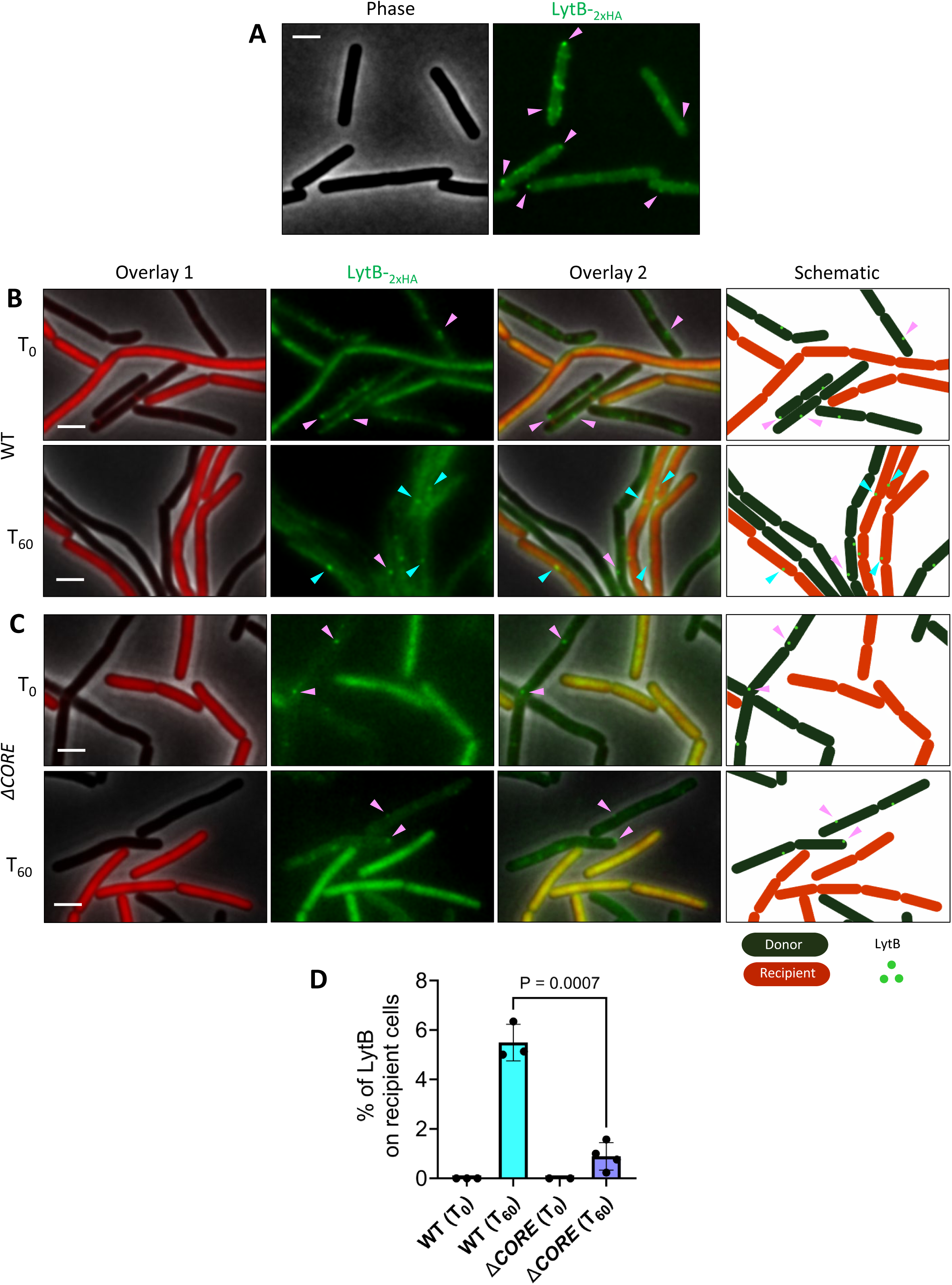
Deposit of LytB molecules on recipient bacteria. **(A)** AB266 (*lytB-_2XHA_*) cells were grown in liquid LB medium and surface-labeled with anti-HA Qdot525 antibodies to follow LytB-_2xHA_ molecules. Shown are phase contrast (gray) and a signal from anti-HA Qdot525 (green). Arrowheads highlight the localization of LytB molecules on the cell surface. Scale bar, 1 μm. **(B)** AB266 (*lytB-_2XHA_*) cells were grown in liquid LB medium, surface-labeled with anti-HA Qdot525 antibodies, mixed with BDR2637 (P*_veg_-mCherry*) recipient cells in 1:1 ratio, and incubated on LB solid medium. Shown are overlay 1 of phase contrast (gray) with fluorescence from mCherry (red), a signal from anti-HA Qdot525 (green), and overlay 2 of phase contrast, mCherry, and anti-HA Qdot525. Images were captured at t=0, (upper panels), and at 60 min (lower panels). Right panels depict schematics of donor cells (black), recipient cells (red), and LytB molecules (green). Pink arrowheads highlight the localization of LytB molecules on donor cells surface, while cyan arrowheads highlight the transferred LytB molecules on recipient cell surface. Scale bars, 1 μm. **(C)** MG85 (Δ*CORE, lytB-_2XHA_*) cells were grown in liquid LB medium, surface-labeled with anti-HA Qdot525 antibodies, mixed with MG21 (Δ*CORE*, P*_veg_-mCherry*) recipient cells in 1:1 ratio, and incubated on LB solid medium. Shown are overlay 1 of phase contrast (gray) with fluorescence from mCherry (red), a signal from anti-HA Qdot525 (green), and overlay 2 of phase contrast, mCherry, and anti-HA Qdot525. Images were captured at t=0, (upper panels), and at 60 min (lower panels). Right panels depict schematics of donor cells (black), recipient cells (red), and LytB molecules (green). Pink arrowheads highlight the localization of LytB molecules only on donor cell surface. Scale bars, 1 μm. **(D)** Quantitation of the experiments described in (B-C). Shown are average number of LytB molecules [%] detected on recipient WT BDR2637 (P*_veg_-mCherry*) (B) or Δ*CORE* MG21 (Δ*CORE*, P*_veg_-mCherry*) (C). LytB molecules were manually counted using NIS analysis software. Shown are mean ± signal from anti-HA Qdot525 (green). *P* values were determined using a one-tailed, unpaired Student’s *t*-test (*p* =0.0007). Results were collected from 3 independent biological repeats for WT (t_0_, *n*_recipient cells_ =405 and t_60_, *n*_recipient cells_ = 1098) and 2 for Δ*CORE* (t_0_, *n*_recipient cells_ =382 and t_60_, *n*_recipient cells_ = 1391).

**Figure S5.**
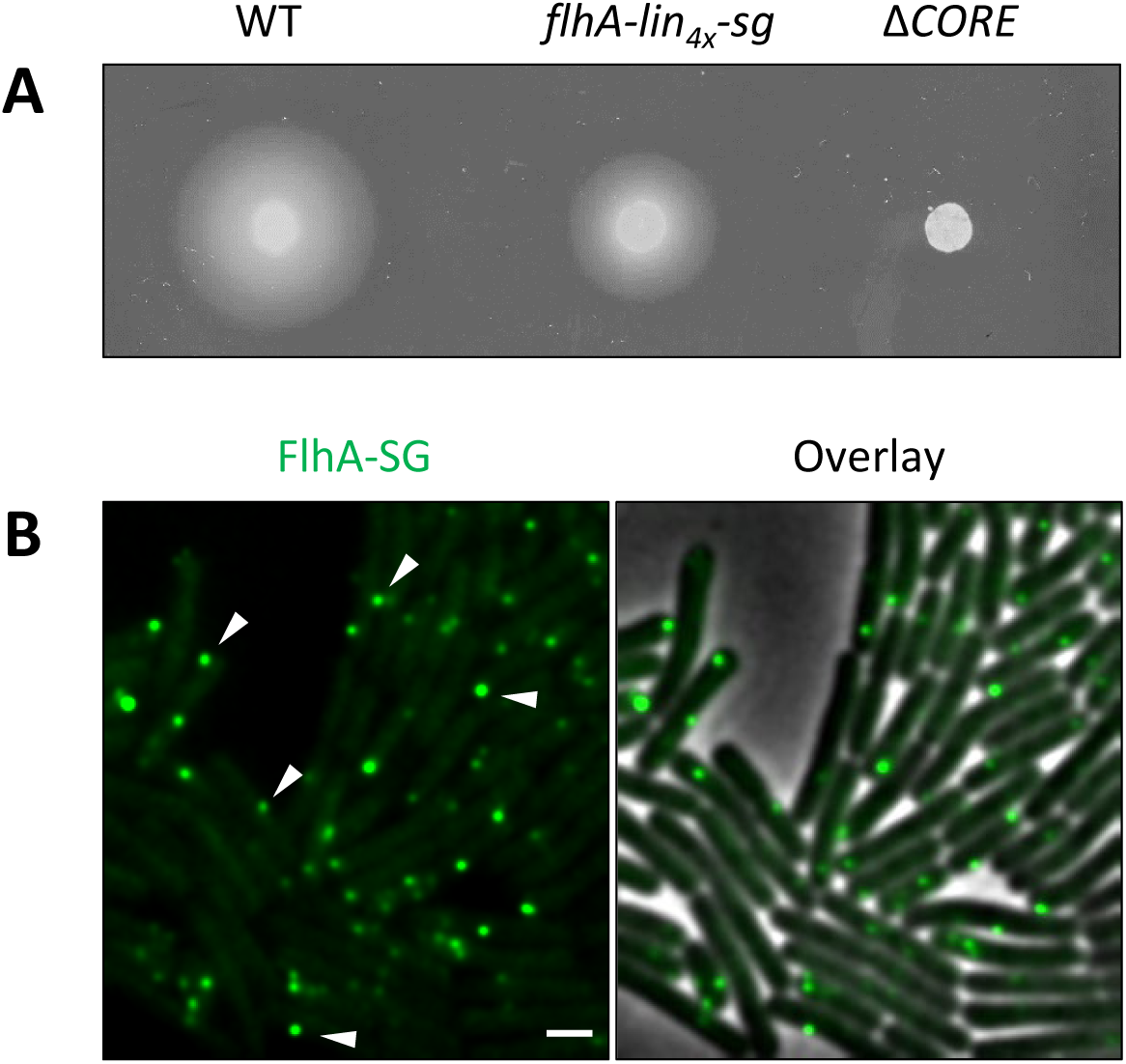
Functionality of FlhA-SG fusion. **(A)** Wild-type (PY79), GV591 (*flhA-lin_4x_-sg*), and SH9 (Δ*CORE*) cells were grown to the mid-logarithmic phase, and motility was examined by spotting the strains onto LB plates containing 0.3% agar. Cells were photographed after 7 h of incubation at 37°C. **(B)** GV591 (*flhA-lin_4x_-sg*) cells were grown in liquid LB medium and observed by fluorescence microscopy. Shown are FlhA-SG (green), and its overlay with phase contrast (gray). Arrowheads highlight FlhA-SG foci. Scale bar, 1 μm.

**Figure S6.**
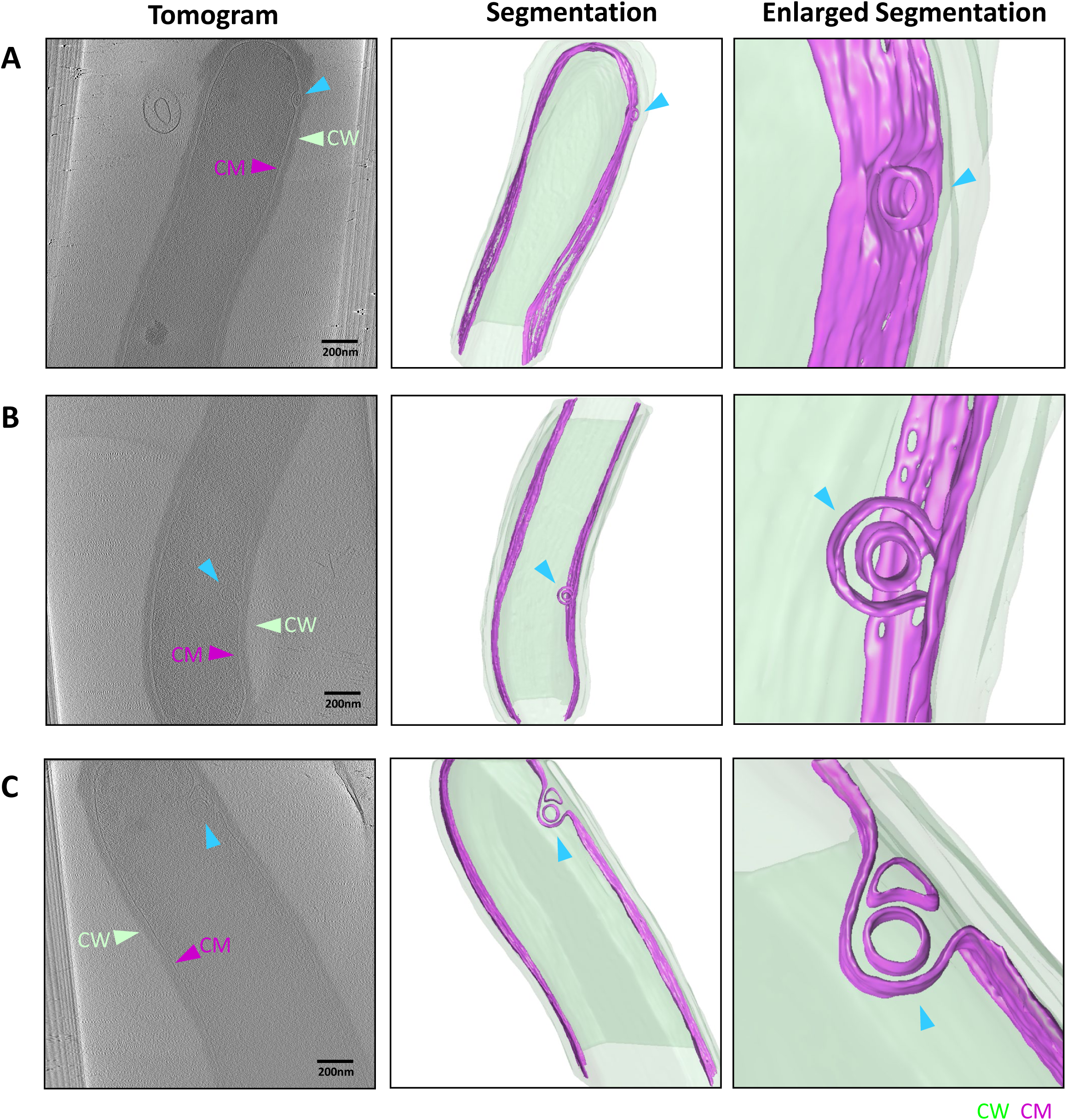
Examples of cells displaying different-sized tubeosomes. **(A-C)** AB328 (Δ*ponA*, P_IPTG_*-ymdB*, Δ*hag*) cells were grown on solid LB medium and subjected to Cryo-ET. Left panels: slices through cryo-electron tomograms displaying cells harboring prominent tubeosomes of different sizes; Middle panels: corresponding 3D segmentations; Right panels: zoomed-in views of tubeosome structures. 3D Segmentations: CW (cell wall; green), CM (cell membrane; purple). Arrowheads highlight CW (cell wall; green), CM (cell membrane; purple), tubeosome (blue).

**Figure S7.**
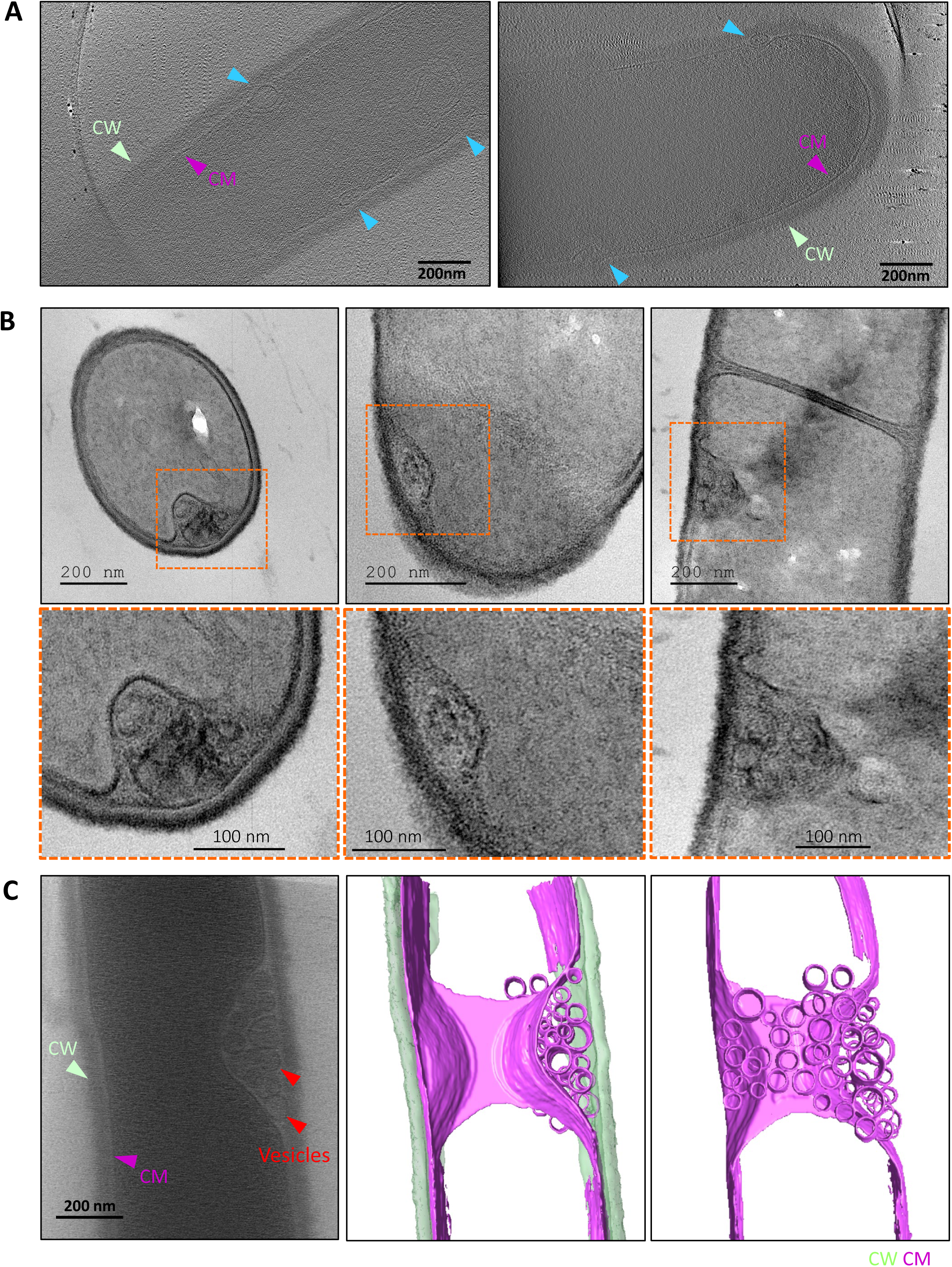
Visualization of membrane invagination by TEM and Cryo-ET. **(A)** AB328 (Δ*ponA*, P_IPTG_*-ymdB*, Δ*hag*) cells were grown on solid LB medium and subjected to Cryo-ET. Shown are slices through cryo-electron tomograms from cells displaying multiple tubeosomes. Arrowheads highlight CW (cell wall; green), CM (cell membrane; purple), tubeosome (blue). **(B)** PY79 (WT) cells were grown in LB to mid-log phase in gentle shaking, fixed, and prepared as longitudinal and transverse ultra-thin sections for visualization by TEM. Shown are cells displaying tubeosome-like structures (upper panels) and their inset enlargement (lower panels). **(C)** MG164 (Δ*ponA*, Δ*CORE*) cells were grown on solid LB medium and subjected to Cryo-ET. Left panel shows a slice through a cryo-electron tomogram of a cell harboring stacks of membrane vesicles, while the middle and right panels show the 3D segmentation of the vesicle compartment from different viewpoints. Arrowheads highlight CW (cell wall; green), CM (cell membrane; purple), tubeosome (blue), and vesicles (red).

## Supplemental Movie Legends

**Movie S1. Networks of Intercellular nanotubes**

GB168 (P_IPTG_*-ymdB*) cells were grown on a solid LB medium containing FM4-64 (red), and time-lapse microscopy was performed. Shown are overlay images of phase contrast (gray) and nanotubes (red, Z_N_) captured at 300 ms/frame (total 83 s). Corresponding to Figure 2A.

**Movie S2. Following the dynamics of an intercellular nanotube**

GB168 (P_IPTG_*-ymdB*) cells were grown on a solid LB medium containing FM4-64 (red), and time-lapse microscopy was performed. Shown are overlay images of phase contrast (gray) and nanotubes (red, Z_N_) captured at 200 ms/frame (total 58 s). Corresponding to Figure 2B.

**Movie S3. Recording extending nanotube dynamics**

MG200 (P_IPTG_*-ymdB,* P1*_rrnB_-gfp*) cells, expressing GFP, were grown on a solid LB medium containing FM4-64 (red), and time-lapse microscopy was performed. Shown are overlay images of cells (green) and nanotubes (red, Z_N_), captured at 300 ms/frame (total 64 s). Corresponding to Figure 2D.

**Movie S4. Visualizing the formation of an intercellular nanotube bridge**

GB168 (P_IPTG_*-ymdB*) cells were grown on a solid LB medium containing FM4-64 (red), and time-lapse microscopy was performed. Shown are overlay images of phase contrast (gray) and nanotubes (red, Z_N_) captured at 200 ms/frame (total 238 s). Corresponding to Figure 3A.

**Movie S5. Nanotube shortening following the formation of an intercellular bridge**

GB168 (P_IPTG_*-ymdB*) cells were grown on a solid LB medium containing FM4-64 (red), and time-lapse microscopy was performed. Shown are nanotubes (Z_N_) captured at 200 ms/frame (total 75 s). Corresponding to Figure 3D.

**Movie S6. Nanotubes protrusion from sites of membrane accumulation**

GB168 (P_IPTG_*-ymdB*) cells were grown on a solid LB medium containing FM4-64 (red), and time-lapse microscopy was performed. Shown are nanotubes (Z_N_) captured at 400 ms/frame (total 235 s). Corresponding to Figure 4B.

**Movies S7-S8. The existence of the tubeosome**

Cryo-electron tomography of AB328 (Δ*ponA*, P_IPTG_*-ymdB*, Δ*hag*) cells and a 3D segmentation thereof, indicating the presence of tubeosome structures. Scale bars, 100 nm. Movie S7 corresponds to Figure 5A-5B and Movie S8 to Figure 5C-5E.

**Movie S9. Extending membranous nanotube**

Cryo-electron tomography of AB328 (Δ*ponA*, P_IPTG_*-ymdB*, Δ*hag*) cell and a 3D segmentation thereof, indicating the presence of an extending nanotube. Scale bars, 100 nm. Corresponding to Figure 6A-6B.

**Movie S10. Δ*CORE* cells exhibit vesicle sacs**

Cryo-electron tomography of MG164 (Δ*ponA*, Δ*CORE*) cell and a 3D segmentation thereof, indicating the presence of multiple stacks of membrane vesicles in the space between the cell membrane and the cell wall. Scale bars, 100 nm. Corresponding to Figure 6C-6D.

## Methods

### Bacterial strains and growth conditions

*B. subtilis* strains used in this study are derivatives of wild-type PY79 ^1^. *B. megaterium* is a wild soil isolate OS2 strain ^2^. Bacterial strains and primers, utilized in this study, are listed in Tables S1 and Table S2, respectively. Antibiotic resistance cassettes flanked by lox66/lox71 sites were PCR-amplified from pWX465 (*cat*), pWX466 (*spec*), pWX467 (*erm*), pWX469 (*tet*), and pWX470 (*kan*) using primers 2430 and 2431. Plasmid constructions were performed in *E. coli* DH5α using standard molecular biology techniques. All general methods were performed as described previously ^3^. Briefly, *B. subtilis* overnight cultures were diluted to 0.05 OD_600nm_ and grown to a density of 0.8 OD_600nm_ in LB medium (BD Biosciences, Difco^TM^) at 37°C. A single colony of *B. megaterium* was grown in liquid LB medium up to 0.8 OD_600nm_. Gene expression from P_IPTG_ was induced with 0.1-0.5 mM IPTG (isopropyl-β-D-1-thiogalactopyranoside; Sigma-Aldrich). When needed, the following antibiotics were used for *B. subtilis* selection: kanamycin (10 µg/ml, US Biological), chloramphenicol (5 µg/ml, Sigma-Aldrich), lincomycin (25 µg/ml, Sigma-Aldrich), erythromycin (1 µg/ml, Sigma-Aldrich), tetracycline (10 µg/ml, Sigma-Aldrich), spectinomycin (100 µg/ml, Sigma-Aldrich), ampicillin 100 µg/ml (Sigma-Aldrich).

### Fluorescence microscopy

For real-time imaging of bacterial nanotubes, a specialized metal chamber (A-7816, ThermoFisher Scientific, USA) was assembled to observe bacterial growth and nanotube dynamics on solid LB medium by fluorescence microscopy. Imaging was performed using ECLIPSE Ti2 microscope (Nikon, Japan) equipped with a Prime BSI camera (Photometrics, USA) in a temperature-controlled incubation chamber (Okolab). To improve visualization of bacterial nanotubes, LB medium was supplemented with UltraPure™ 1.5% agarose (Invitrogen, USA) and 0.5% Gellan Gum (GEL-GRO^TM^, ICN Biomedicals, Inc., USA or MP Biomedicals, USA). The mixture was melted and immediately poured into the metal chamber to form a firm and transparent agarose pad, as schematically shown in Figure 1A. To visualize cells and nanotubes, FM4-64 (ThermoFisher Scientific, USA) fluorescent membrane dye was added to the medium at a final concentration of 10 μg/ml. Cells were incubated at 30-32°C, followed by time-lapse fluorescence microscopy using 100x objective. To observe the localization of FlhA-StayGold, exponentially growing cells were harvested at 0.8 OD_600nm_, spotted onto an agarose pad, and imaged. System control and image analysis were performed using NIS-Elements AR analysis (version 5.30.07, Nikon, Japan) and Fiji (Image J). When needed, captured images were 2D and 3D deconvolved with Richardson–Lucy iterations methods, using NIS-Elements AR analysis software (version 5.30.07, Nikon, Japan).

### Live immunofluorescence microscopy

*B. subtilis* cells were grown until 0.8 OD_600nm_ at 37°C in a roller. Cells (1 ml) were harvested by centrifugation (2400 rpm, 6 minutes), and the resulting pellet was washed by resuspension in fresh LB medium. For immunostaining, cells were incubated with anti-HA primary antibodies (1:200 in LB; Invitrogen, USA) for 60 minutes, followed by three washes with LB. Subsequently, cells were incubated with Qdot™ 525-conjugated secondary antibodies (1:700 in LB; Invitrogen, USA; anti-rabbit IgG). After additional washes with LB, cells were stained with FM4-64 fluorescent membrane dye, spotted onto an LB agarose pad, and visualized by fluorescence microscopy using 100 x objective. For assessing LytB molecule transfer, donor and recipient cells were grown separately in LB medium to 0.8 OD_600nm_ at 37°C. Donor cells were labeled with antibodies as described above, while recipient cells remained unlabeled. Donor and recipient cells were then mixed at a 1:1 ratio, spotted onto LB agarose pad, and images were captured at the indicated time points.

### Analysis of intercellular nanotube movement amplitude

Nanotube movement amplitude during dynamic movement was generated in Fiji (ImageJ) by computing the sum projection from of all 229 recorded frames shown in Figure 2B. Fluorescence intensity of nanotube for three selected regions (cyan, yellow and green) was measured in both static and dynamic states using the Fiji (ImageJ) ‘Profile Line’ tool.

### Tracking of nanotube dynamics

Tracking analysis was performed using the TrackMate ^4^ plugin in Fiji (ImageJ). Trajectories were generated for each nanotube movement, and displacement and velocity were subsequently calculated from the resulting tracking data.

### Motility assay

Motility assay was performed as described ^5^ with some modifications. Briefly, cells were grown to the mid-logarithmic phase in LB and then concentrated 10-fold to an OD_600nm_ 0.5. Cell suspension (5 μl) was gently spotted on freshly prepared 0.3% LB agar plates. The plates were incubated for 7 hours at 37°C and imaged using a photo scanner (EPSON-V800).

### Ultramicrotome sections and transmission electron microscopy

*B. subtilis* cells were grown in liquid LB with slow shaking, gently centrifuged, and samples were fixed in 2% formaldehyde and 2.5% glutaraldehyde in 0.1 M sodium cacodylate buffer, pH 7.4 with post-fixation in 2% OsO_4._ Samples were processed for transmission electron microscopy, following embedding in Agar 100 resin (Agar Scientific, UK). Thin sections (70-90 nm) were prepared using an Ultracut UCT microtome (Leica), stained with Uranyl acetate and Lead citrate. Grids were viewed with Jeol® JEM-1400 Plus TEM (Jeol®, Tokyo, Japan), equipped with ORIUS SC600 CCD camera (Gatan®, Abingdon, United Kingdom), and Gatan Microscopy Suite program (Digital Micrograph, Gatan®, UK).

### Bacterial sample preparation for cryo-ET and cryo-CLEM

*B. subtilis* cells were grown until 0.8 OD_600nm_ (37°C, LB) with slow shaking (25 rpm). Cells were harvested by gentle centrifugation at 4000 rpm for 5 min. For cryo-CLEM experiments, cells were stained with FM4-64 (10 μg/ml) for 1 minute at room temperature. Subsequently, 4-5 μL of the sample were spotted on Quantifoil gold grids (Electron Microscopy sciences, USA), positioned on top of LB agar plates, and incubated for 1-2 hrs at 37°C to prepare cryo-ET samples.

### Sample preparation for cryo-ET and cryo-CLEM

Quantifoil 200 mesh R2/2 Au (LF) grids (Electron Microscopy sciences, USA) were glow-discharged by Gatan Solarus model 950 system (30 seconds, 20 watts), positioned on top of LB agar plates, and cells were grown on the grids for 1 hr at 37°C. Grids were picked up with a tweezer and gently washed with Phosphate-Buffered Saline (PBSx1, pH 7.4) to remove loose layers of cells before loading to Vitrobot Mark IV System (ThermoFisher Scientific, USA). The Vitrobot chamber was maintained in a 100% humidity at 22°C. Once in the Vitrobot chamber, 4 μL of bovine serum albumin-treated 10 nm gold bead solution was loaded onto the grid, which was subsequently blotted to remove excess liquid and plunge-frozen in liquid ethane.

### Cryo-focused ion beam milling of bacterial cell clusters

Samples were thinned by focused ion beam milling, which was performed on an Aquilos 2 cryo-DualBeam instrument (ThermoFisher Scientific, USA). Subsequent to plunge freezing, grids were clipped into standard CryoFIB AutoGrid Rings (ThermoFisher Scientific, USA) with a cutout to allow for shallower milling angles and were marked in order to monitor the milling direction during cryo-ET data acquisition. Samples were coated in-column with platinum [20 s, 20.0 mA, 10 Pa] and then sputtered with a layer of organometallic platinum using the gas injection system GIS [35s]. Milling included rough, medium and fine milling, with respective electric current of 0.3 nA, 0.1 nA and 0.05 nA, and was followed by two polishing steps, with electric current values equal to 30 pA and 10 pA. The preparation of lamellae was stepwise, with target thickness of ∼200 nm. Samples were stored in liquid nitrogen until cryo-ET imaging with Titan Krios was performed.

### Cryo- CLEM and cryo-ET data collection and processing

For cryo-CLEM experiments, plunge-frozen samples were examined with STELLARIS cryo-confocal system to identify targets suitable for cryo-ET data collection. The system is part of the Advanced Electron Microscopy Facility at University of Chicago. Cryo-ET data collection was performed on a Titan Krios G3i 300 keV field electron microscope (ThermoFisher Scientific, USA) that equipped with a Gatan K3 camera and a BioQuantum energy filter operated using either Tomo5 software or SerialEM ^6^. Tilt series were collected either using dose symmetric tilt-scheme ^7^ from -51° to +51° (at 3° tilt increments) at -7.00 µm defocus under low dose conditions at a pixel size of 3.34 Å, or using fast-incremental tilt series scheme ^8^. A cumulative dose of ∼120 e−/Å^2^ was used. Cryo-ET data was collected at Advanced Electron Microscopy Facility at the University of Chicago and the Cryo-EM facility at Caltech.

### Cryo-ET data processing and visualization

Three-dimensional reconstructions of tilt series were performed either using the IMOD software package ^9^, or automatically with *Tomo Live* software ^10^ at the University of Chicago and the RAPTOR pipeline ^11^ at Caltech. Tomogram segmentations were performed using the Dragonfly software packages (https://www.theobjects.com/dragonfly/index.html). A series of TIFF files were generated from an MRC tomogram using “mrc2tif” command in the IMOD software package. The generated TIFF image stack was imported into Dragonfly followed by intensity scale calibration. Initial training models were created in the Segmentation Wizard for deep learning. At least three representative tomogram slices were selected to train each model. Regions of interest (ROIs), such as membrane and cell wall, were labeled as different color classes, and the remainder of the unlabeled region was recognized as background. The AI-driven algorithm constructs 3D models through tracing the 2D gray contrast of each tomogram slice. The generated model was then modified manually to remove noise and fill in the missing parts. The Smooth Mesh tool was applied for polishing the model surface. Dragonfly Movie Maker was used to generate the movies.

### Quantification and statistical analysis

All statistical analyses, data processing, and figure preparation were performed using GraphPad Prism software. Quantifications of tubeosome structures were conducted manually, as described in the respective figure legends. All microscopy image quantifications were performed using multiple biological replicates, as indicated in the corresponding figure legends.

---

**Table S1:**
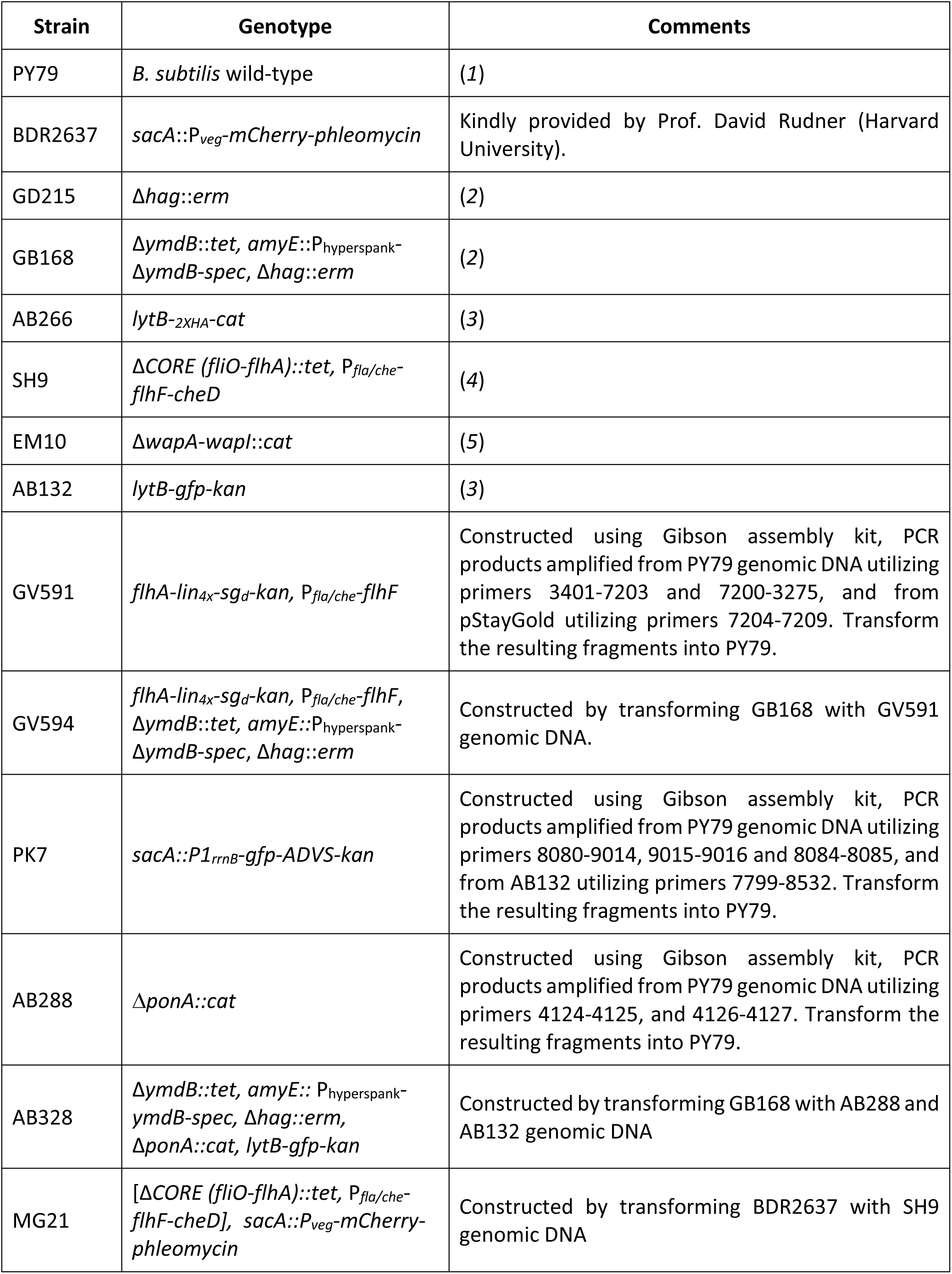
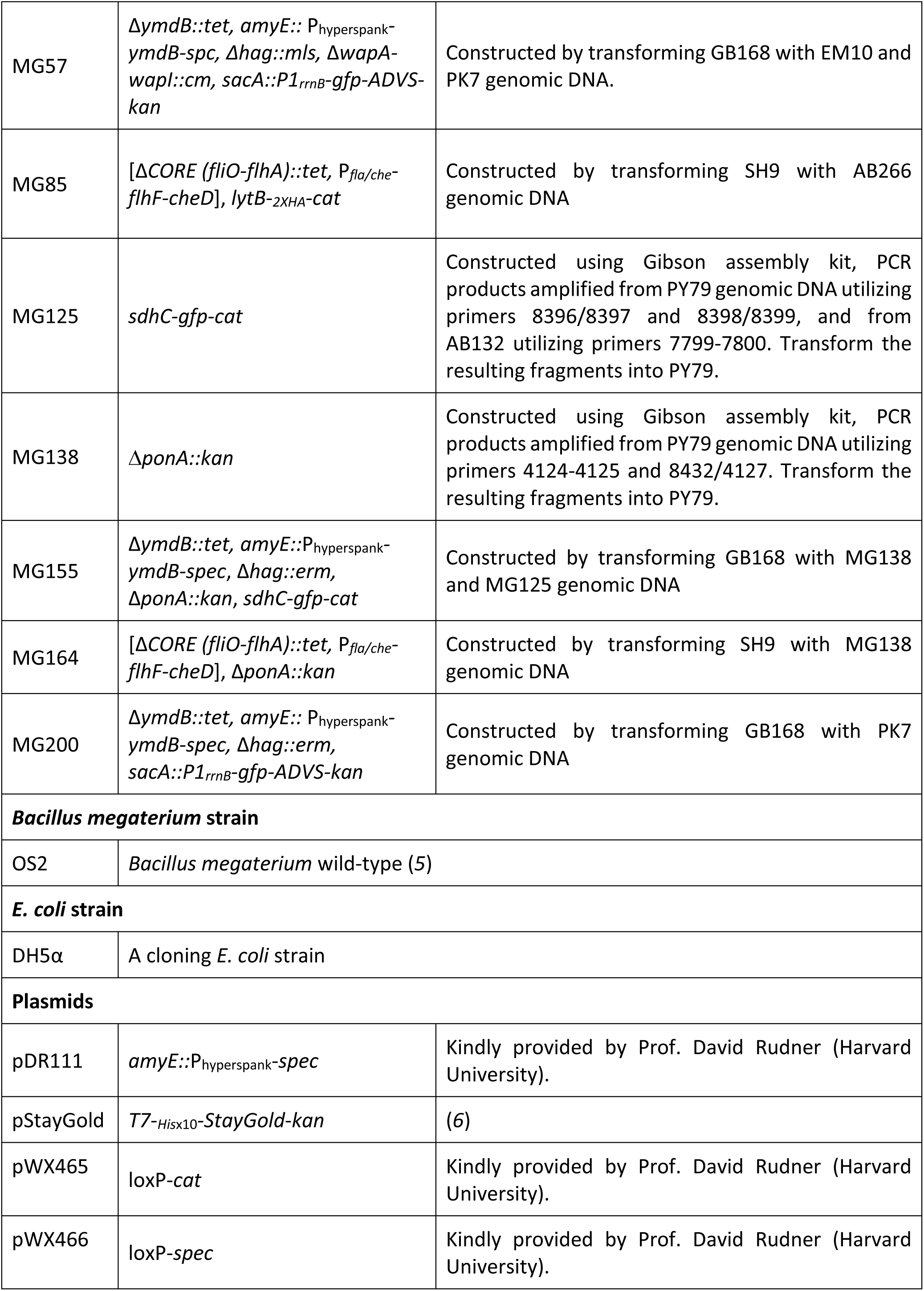
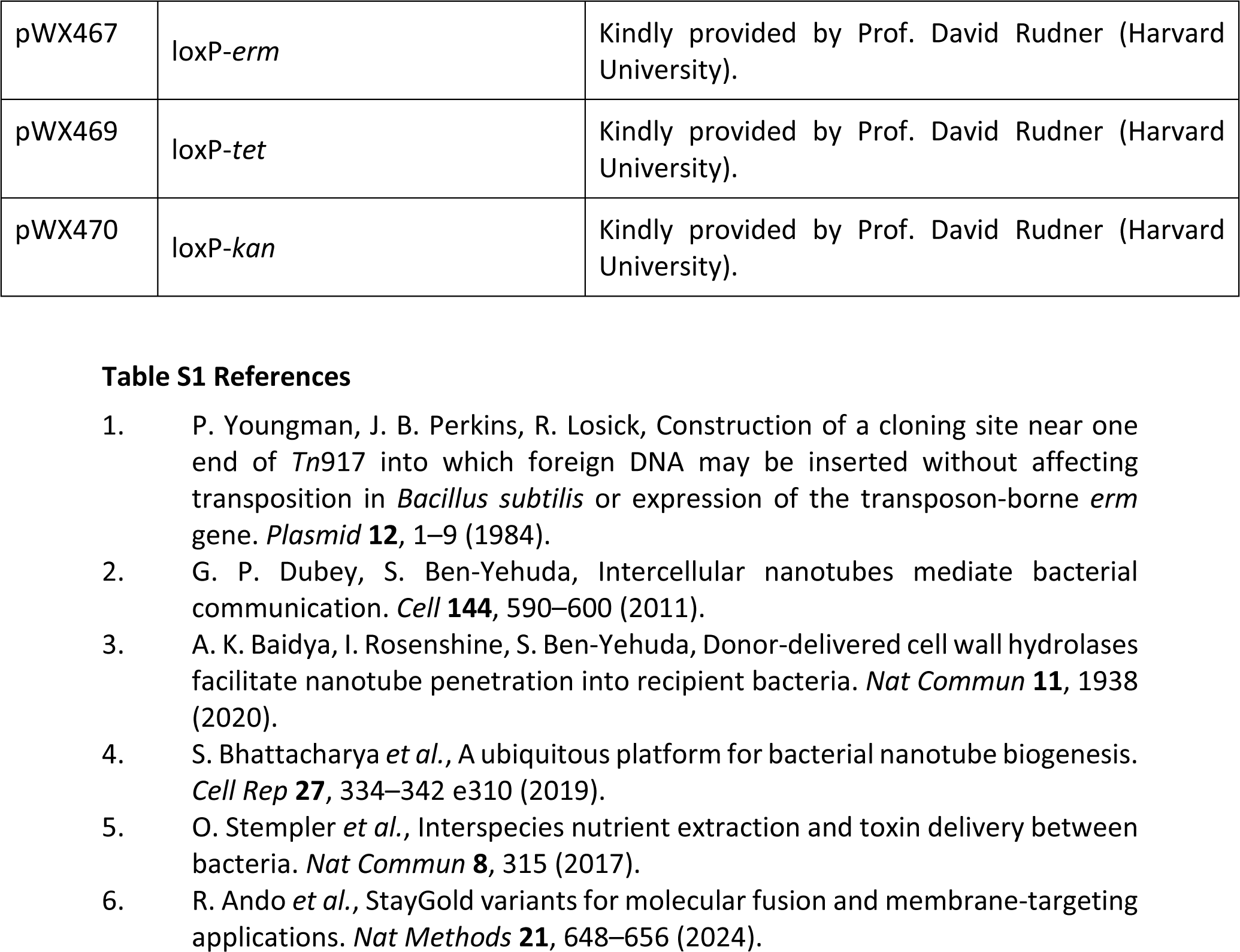
Strains and plasmids used in this study.

**Table S2:**
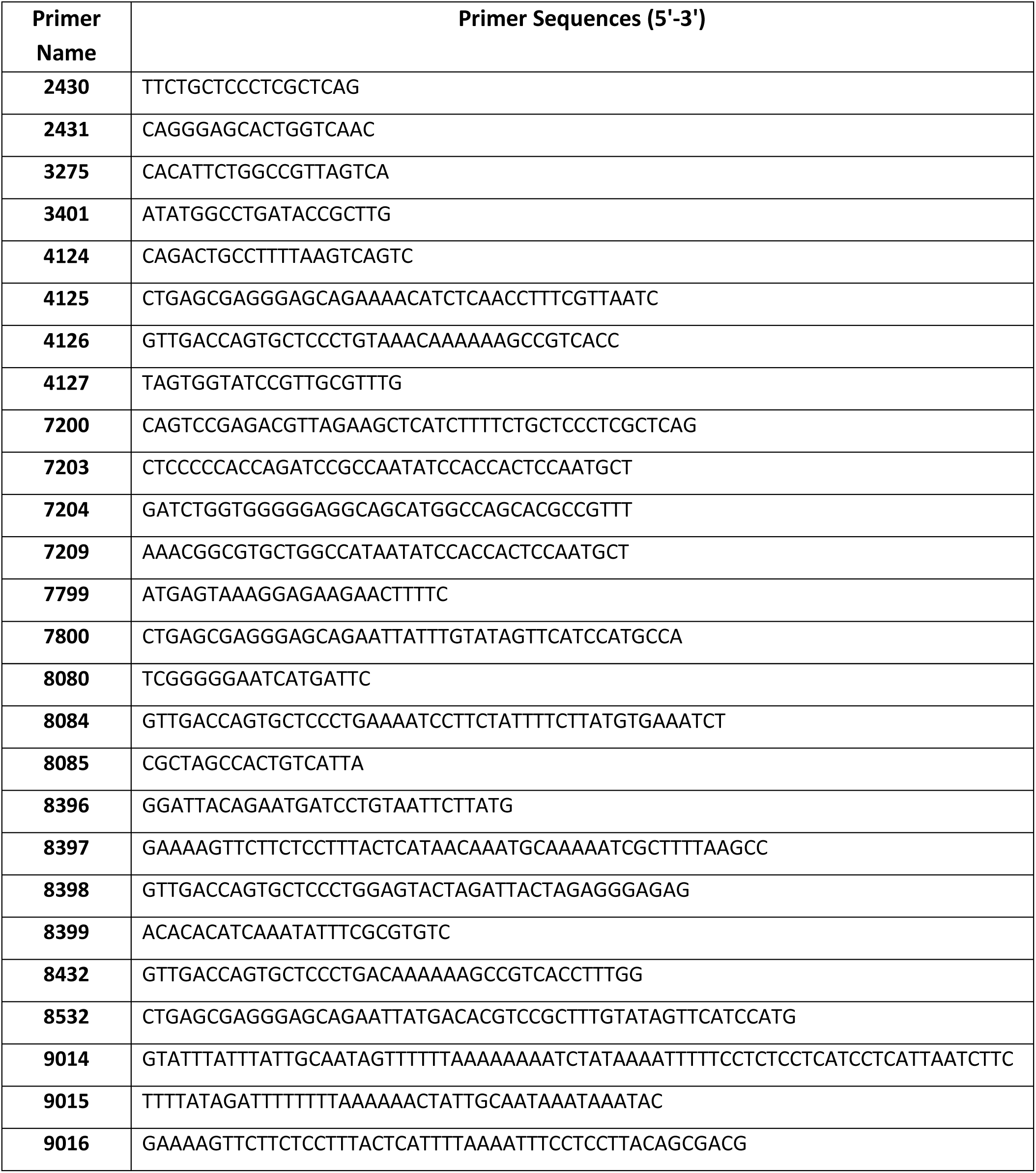
List of primers used in this study.

